# Conserved Transcription Factors Control Chromatin Accessibility and Gene Expression to Maintain Cell Fate Stability and Restrict Reprogramming of Differentiated Cells

**DOI:** 10.1101/2021.06.28.450261

**Authors:** Maria A. Missinato, Sean A. Murphy, Michaela Lynott, Anaïs Kervadec, Michael S. Yu, Yu-Ling Chang, Suraj Kannan, Mafalda Loreti, Christopher Lee, Prashila Amatya, Hiroshi Tanaka, Chun-Teng Huang, Pier Lorenzo Puri, Chulan Kwon, Peter D. Adams, Li Qian, Alessandra Sacco, Peter Andersen, Alexandre R. Colas

**Author notes:** Equal Contribution (Maria A. Missinato, Sean A. Murphy).

## Abstract

The comprehensive characterization of mechanisms safeguarding cell fate identity in differentiated cells is crucial for *1)* our understanding of how differentiation is maintained in healthy tissues or misregulated in disease states and *2)* to improve our ability to use direct reprogramming for regenerative purposes. To uncover novel fate-stabilizing regulators, we employed a genome-wide TF siRNA screen followed by a high-complexity combinatorial evaluation of top performing hits, in a cardiac reprogramming assay in mouse embryonic fibroblasts, and subsequently validated our findings in cardiac, neuronal and iPSCs reprogramming assays in primary human fibroblasts and adult endothelial cells. This approach identified a conserved set of 4 TFs (ATF7IP, JUNB, SP7, and ZNF207 [AJSZ]) that robustly opposes cell fate reprogramming, as demonstrated by up to 6-fold increases in efficiency upon AJSZ knockdown in both lineage- and cell type-independent manners. Mechanistically, ChIP-seq and single-cell ATAC-seq analyses, revealed that AJSZ bind to both open and closed chromatin in a genome-wide and regionalized fashion, thereby limiting reprogramming TFs access to target DNA and ability to remodel the chromatin. In parallel, integration of ChIP-seq and RNA-seq data followed by systematic functional gene testing, identified that AJSZ also promote cell fate stability by proximally down-regulating a conserved set of genes involved in the regulation of cell fate specification (MEF2C), proteome remodeling (TPP1, PPIC), ATP homeostasis (EFHD1), and inflammation signaling (IL7R), thereby limiting cells ability to undergo large-scale phenotypic changes. Finally, simultaneous knock-down of AJSZ in combination with cardiac reprogramming TFs overexpression improved heart function by 250% as compared to no treatment and 50% as compared to MGT, 1 month after myocardial infarction. In sum, this study uncovers a novel evolutionarily conserved mechanism mediating cell fate stability in differentiated cells and also identifies AJSZ as promising therapeutic targets for regenerative purposes in adult organs.

**Significance Statement:** Differentiated cells can be converted from one cell type into another by overexpressing lineage-determining transcription factors. Direct lineage reprogramming represents a promising strategy for regenerative medicine, but current clinical applications remain limited by the low yield of the reprogramming process. Here, we present the identification and detailed mechanistic study of a novel mechanisms opposing reprogramming process and promoting cell fate stability. Using a highthroughput screening technique, we identified four transcription factors that act as blockades to cell type change. By tracking chromatin and gene expression changes, we reconstructed a conserved pathway mediating cell fate stability in differentiated cells.

## Introduction

Cell fate identity is acquired during the process of differentiation and consists in the establishment of a lineage-specific combination of chromatin architecture and gene expression^1^, that supports the function of specialized cells in multi-cellular organisms. Although observed to be stable in differentiated cells, groundbreaking studies^2–4^ have revealed that cell fate identity can be reprogrammed by overexpressing lineage-specific combinations of transcription factors (TFs)(reviewed in ^5^). In this context, different cell types such as fibroblasts or endothelial cells, could be turned into induced pluripotent stem cells^4,6^, neurons^7^ or cardiomyocytes^8^, however, in most cases only a small fraction of fate-challenged cells underwent reprogramming (reviewed in ^9–11^), thus revealing the existence of robust fate-stabilizing mechanisms opposing this process in differentiated cells, which currently limit our ability to use cell fate reprogramming for regenerative purposes^12,13^.

Intense research (reviewed in ^5,11,14^) during the past decade has established that a rate-limiting step for TF-induced cell fate conversion resides in the ability of reprogramming TFs to efficiently bind to their target DNA, and in turn activate destination cell type gene expression^15^. Consistent with this reprogramming paradigm, epigenetic regulators of heterochromatin formation such as histone chaperones^16^, enzymes involved in histone H3^17,18^ or DNA^19,20^ methylation, were found to oppose cell fate reprogramming *via* their ability to limit reprogramming TFs to access their target DNA. In addition to these chromatin-associated mechanisms, regulators of source and destination cell type transcriptomes such as TGF-β and inflammatory signaling pathways^21,22^ or genes involved in RNA methylation^23,24^, alternative polyadenylation^25^ and splicing^26^, were also found to control cells ability to reprogram by modulating mRNA stability or splicing patterns of genes required for the fate conversion process. Collectively, these studies demonstrate that cell fate identity is actively maintained by the concomitant regulation of cell type-appropriate chromatin architecture and gene expression, however, in this context, it remains to be established whether yet to be discovered mechanism(s) integrating both regulatory dimensions might control this process in differentiated cells.

TFs are essential determinants of differentiation during embryogenesis^27,28^ and mediate their role by direct DNA binding to regulate transcription and chromatin accessibility^29,30^ and thus, represent an ideal class of proteins to promote cell fate stability in differentiated cells. Consistent with our hypothesis, recent work from Gurdon and colleagues^31^, have shown that long-term DNA association of lineage-specific TF, ASCL1, contributes to maintain gene expression and stabilize fate commitment in differentiating cells. Moreover, several studies have also shown that various TFs, including SNAI1^32^, cJun^33^, Bright/ARID3A^34^, oppose cell fate reprogramming by enhancing source cell gene expression or repressing reprogramming TFs-associated gene expression. However, a systematic evaluation of TFs as fate stabilizers has not been conducted and in this context, we will ask whether the function of such regulators might be evolutionarily conserved, lineage and/or cell type-specific or independent. Mechanistically, we will delineate how pre-existing fate-stabilizing TF binding to the DNA might concomitantly orchestrate chromatin accessibility dynamics and transcription to oppose the reprogramming process. Finally, we will test whether the depletion of such fate stabilizers might enhance our ability to use cell fate reprogramming for regenerative purposes *in vivo*.

To uncover novel fate-stabilizing regulators, we employed a genome-wide TF siRNA screen followed by a high-complexity combinatorial evaluation of top performing hits, in a cardiac reprogramming (CR) assay in mouse embryonic fibroblasts, and subsequently validated our findings in cardiac, neuronal and iPSCs reprogramming assays in primary human fibroblasts and adult endothelial cells. This approach identified a conserved set of 4 TFs (ATF7IP, JUNB, SP7, and ZNF207 [AJSZ]) that robustly opposes cell fate reprogramming, as demonstrated by up to 6-fold increases in efficiency upon AJSZ knockdown in both lineage- and cell type-independent manners. Mechanistically, ChIP-seq and single-cell ATAC-seq analyses, revealed that AJSZ bind to both open and closed chromatin in a genome-wide and regionalized fashion, thereby limiting reprogramming TFs access to target DNA and ability to remodel the chromatin. In parallel, integration of ChIP-seq and RNA-seq data followed by systematic functional gene testing, identified that AJSZ also promote cell fate stability by proximally downregulating a conserved set of genes involved in the regulation of cell fate specification (MEF2C), proteome remodeling (TPP1, PPIC), ATP homeostasis (EFHD1), and inflammation signaling (IL7R), thereby limiting cells ability to undergo large-scale phenotypic changes. Finally, lentivirus mediated KD of AJSZ in combination with MGT overexpression improved cardiac function by <u>50%</u> as compared to MGT alone and <u>reduced fibrosis</u>, 1 month after myocardial infarction. Collectively, this study uncovers a novel and general mechanism by which conserved TFs promote cell fate stability by concomitantly restricting chromatin accessibility and transcription of genes required for large-scale phenotypic changes.

## Results

### Genome-wide TF screen identifies novel regulators of cell fate stability in mouse fibroblasts

To identify novel fate stabilizers, we employed an immortalized mouse embryonic fibroblasts (iMGT-Myh6-eGFP-MEFs, referred to hereafter as iMGT-MEFs) that carries a cardiac lineage reporter transgene (*Myh6-eGFP*) and can be directly reprogrammed into cardiomyocyte-like cells (iCMs) upon doxycycline (Dox)-induced overexpression of cardiac reprogramming (CR) TFs *<u>M</u>ef2c, <u>G</u>ata4,* and *<u>T</u>bx5* (MGT)(^35^ and Fig. 1A). Upon doxycycline (1μg/ml) treatment, only ~6% of fate-challenged cells express *Myh6* -eGFP by day 3 (Supplementary Fig. 1A), thus suggesting that under these control conditions, most cells fail to undergo reprogramming, thereby providing an ideal model system for the identification of mechanisms opposing cell fate conversion.

**Fig. 1.**
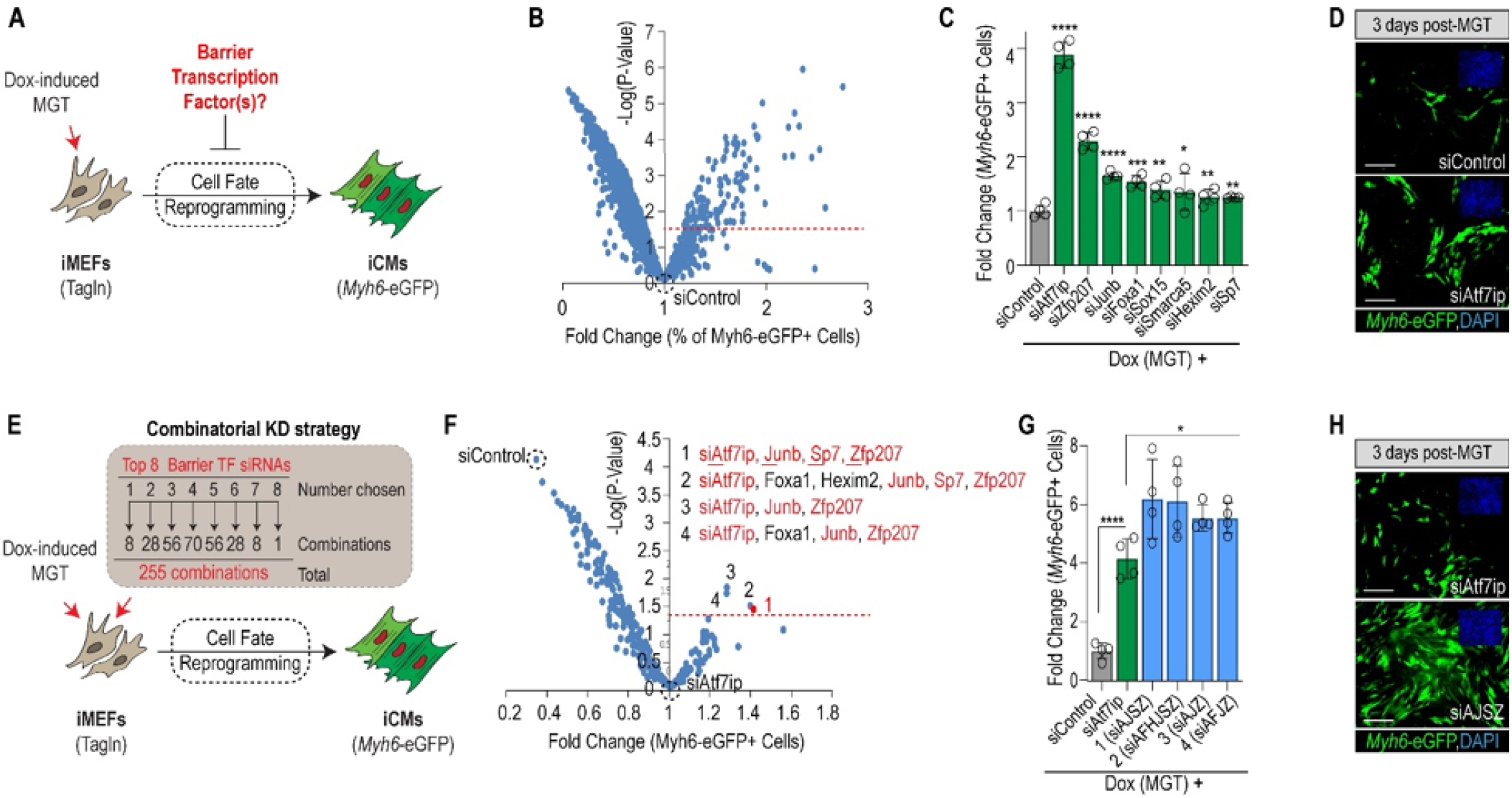
Genome-wide TF screen identifies Atf7ip, JunB, Sp7, and Zfp207 (ZNF207) as barriers to cell fate reprogramming. **(A)** Schematic of the cardiac reprogramming assay and experimental rationale. **(B)** Volcano plot depicting genome-wide TF screen results. X axis shows % of Myh6-EGFP+ positive normalized to siControl. Y axis represents −log of P-value as compared to siControl. The screen was run in experimental quadruplicate. **(C)** Validation of top 20 hits with different siRNA sequences, identifies 8 siRNAs with confirmed activity. Student’s t-test: *p<0.05, **p<0.01, ***p<0.001, ****p<0.0001. **(D)** Representative images of top hit siAtf7ip- and siControl-transfected iMGT-MEFs on day 3 after MGT induction. Myh6-eGFP is shown in green and nuclei are stained with DAPI (blue, top right insets). **(E)** Schematic of combinatorial screening strategy. A total of 255 combinations was tested, each in quadruplicate. (**F)** Volcano plot depicting genome-wide combinatorial screen results. Data were normalized and compared to most potent single hit (siAtf7ip). 1–4 indicate top combinations that were significantly more potent than siAtf7ip alone (p<0.05). si<u>A</u>tf7ip, si<u>J</u>unb, si<u>S</u>p7, and si<u>Z</u>fp207 (siAJSZ) are shown in red. **(G)** Visualization of top 4 siRNA combinations in a histogram plot as in **(C). (H)** Representative images of iMGT-MEFs transfected with siAJSZ (siAtf7ip, siJunb, siSp7, siZfp207) 3 days after MGT induction.

To evaluate the role of TFs as fate stabilizers, we transfected iMGT-MEFs with a library of siRNAs directed against 1435 TFs, one day prior to Dox treatment and quantified CR efficiency (= percentage of Myh6-eGFP+ cells) using automated fluorescence microscopy^36,37^ by day 3. In total, the screen identified 69 siRNAs that significantly increased CR efficiency (>1.25-fold increase in iCMs, p<0.05) as compared to siControl (Fig. 1B and Supplementary Table 1). Next, top 20 hits were tested for validation with independent siRNAs in the iMGT-MEFs assay, which identified 8 distinct siRNAs targeting Atf7ip, Foxa1, Hexim2, Junb, Smarca5, Sox15, Sp7, and Zfp207, that increased CR efficiency (1.2–3.8-fold, p<0.05) as compared to siControl (Fig. 1C, D and Supplementary Fig. 1B), and thus identifying these TFs as novel barriers to CR promoting cell fate stability in iMEFs.

To determine whether these TFs might functionally interact to mediate their fate-stabilizing role, we assembled and tested all possible combinations of the 8 validated siRNAs (255 combinations in total) for function in the iMGT-MEFs assay (Fig. 1E). This approach identified 4 siRNA combinations that significantly enhanced CR efficiency beyond best single siRNA: siAtf7ip (Fig. 1F and Supplementary Table 2). Remarkably, most efficient combination, consisting of siRNAs directed against Atf7ip, Junb, Sp7, and Zfp207 increased the percentage of Myh6-eGFP+ cells from ~6% to ~36%, representing a ~6-fold increase in CR efficiency as compared to siControl and ~1.5-fold increase as compared to siAtf7ip alone (Fig. 1G, H). Taken together, our results identify <u>A</u>tf7ip, <u>J</u>unb, <u>S</u>p7, and <u>Z</u>fp207 (hereafter referred to as A, J, S, and Z), which are members of the ATF-interacting, AP-1, Sp, and Zinc Finger protein families respectively, as novel and functionally interacting barriers to cell fate reprogramming mediating cell fate stability in iMEFs.

### The fate-stabilizing role of AJSZ is conserved in human primary fibroblasts

To determine whether the fate-stabilizing function of AJSZ is evolutionarily conserved, we examined their role during CR in human dermal fibroblasts (HDFs). We first confirmed that AJSZ are expressed in HDFs (Fig. 2A) and that siAJSZ transfection could effectively suppress AJSZ expression (Supplementary Fig. 2A). Next, HDFs were transfected with either siAJSZ or siControl, one day prior to retrovirus-mediated MGT overexpression and effect on CR efficiency was first assessed by qRT-PCR for cardiac-specific markers at day 3. Remarkably, this analysis revealed a significant increase in cardiac-specific gene expression (2–8-fold, p<0.05), including sarcomeric transcripts (*ACTC1, MYL7, TNNT2*), ion channels (*SCN5A, RYR2*), and cardiokines (*NPPA, NPPB*), in siAJSZ-transfected compared with siControl-transfected HDFs (Supplementary Fig. 2B). Consistent with these findings, immunostaining for cardiac marker α-actinin (ACTN1) by day 30 after MGT overexpression, revealed a ~3.2-fold increase in the generation of ACTN1+ cells (from 5% to 16%) in siAJSZ condition as compared to siControl (p<0.05; Fig. 2B, C). Importantly, the increase in CR efficiency observed after AJSZ KD was specifically accompanied by the acquisition of mature structural and functional cardiac phenotypes, including sarcomeric-like structures and calcium transients (Supplementary Fig. 2C, D and Supplementary Movie 1). Collectively, these results indicate that the fate stabilizing function of AJSZ is conserved between mouse and human fibroblasts.

**Fig. 2.**
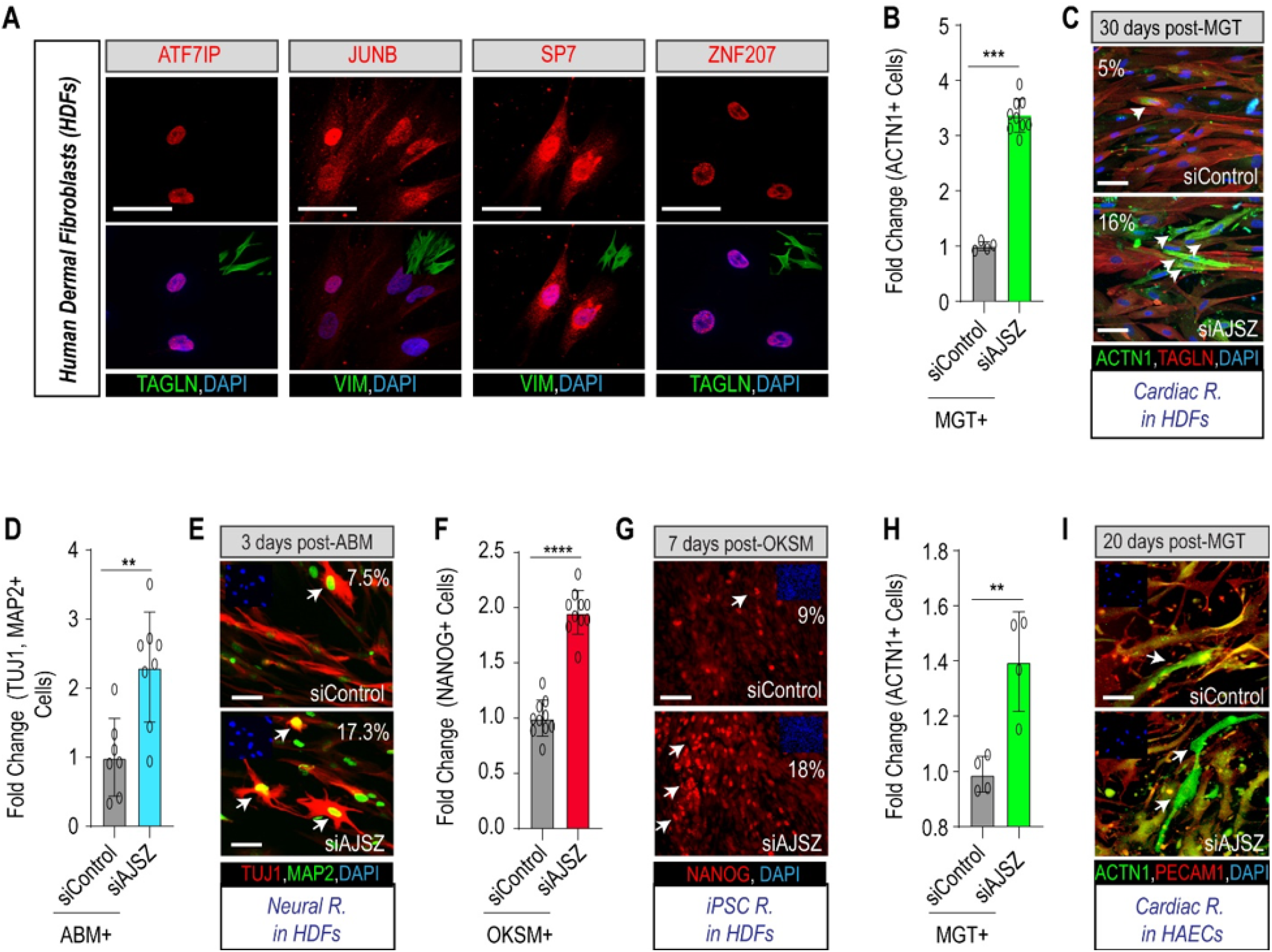
Lineage- and cell type-independent role for AJSZ as barriers to reprogramming. **(A)** Expression of AJSZ in HDFs. Immunostaining for AJSZ is shown in red and fibroblasts markers Vimentin (VIM) or Transgelin (TAGLN) are shown in green. Nuclei are stained with DAPI. **(B)** Quantification of the % of cardiac marker ACTN1-expressing cells normalized to MGT+ siControl condition after 30 days of cardiac reprogramming in HDFs. **(C)** Representative images of HDFs treated with MGT+siControl or MGT+siAJSZ and immunostained for cardiac (ACTN1, green) and fibroblast (TAGLN, red) markers. Nuclei are stained with DAPI (blue). White arrows indicate iCMs. **(D)** Quantification of the % of MAP2 (green) and TUJ1 (red)-double positive cells, 3 days after overexpression of Ascl1, Brn2 and Mytl1 (ABM) in HDFs. **(E)** Representative images of HDFs treated with ABM+siControl or ABM+siAJSZ and immunostained for MAP2 (green) and TUJ1 (red). Nuclei are stained with DAPI (blue, top left insets). White arrows indicate TUJ1+ MAP2+ double-positive cells= iNeurons. **(F)** Quantification of the % of NANOG-positive cells, 7 days after overexpression of OCT4, KLF4, SOX2 and cMYC (OKSM) in HDFs. **(G)** Representative images of HDFs treated with OKSM+siControl or OKSM+siAJSZ and immunostained for pluripotent marker NANOG (red). Nuclei are stained with DAPI (blue, top left insets). White arrows indicate NANOG-positive cells= iPSCs. **(H)** Quantification of the % of ACTN1+ 20 days after MGT overexpression. **(I)** Representative images of HAECs treated with MGT+siControl or MGT+siAJSZ and immunostained for the endothelial marker (PECAM1, red) and the cardiac marker (ACTN1, green). Nuclei are stained with DAPI (blue). White arrows indicate iCMs. Scale bars: 50 μm. Student’s t-test: *p<0.05, **p<0.01, ***p<0.001, ****p<0.0001.

### AJSZ regulate cell fate stability in a lineage- and cell type--independent manner

Next, to assess whether the fate-stabilizing role of AJSZ was restricted to the cardiac lineage or whether it could be generalized to other lineages, we tested their role during the process of neural reprogramming by over-expressing, <u>A</u>scl1, <u>B</u>rn2 and <u>M</u>ytl1 (ABM)38 in HDFs (Supplementary Fig. 3A). Remarkably, compared with siControl-transfected cells, AJSZ KD enhanced by ~2.3-fold (from 7.3% to ~17%) the generation of MAP2+ and TUJ1+ neuron-like cells by day 3 (Fig. 2D, E) and concomitantly increased expression of neuron-specific markers39, including *vGLUT2, GAD67, PVALB,* and *SYN1* by ~2-fold (p <0.05) (Supplementary Fig. 3B), thus suggesting that the AJSZ-mediated regulation of cell fate stability in HDFs is lineage-independent.

To further expand the scope of our findings, we next evaluated whether the role of AJSZ was restricted to direct reprogramming processes^40^ or whether they could be generalized to iPSC reprogramming^4^. To this end, siControl- and siAJSZ-transfected HDFs were transduced with lentiviruses overexpressing pluripotency TFs, *<u>O</u>CT4, <u>K</u>LF4, <u>S</u>OX2* and *c<u>M</u>YC* (OKSM)^41^ and cultured for 7 days. Remarkably, AJSZ KD increased the generation of NANOG+ cells (Fig. 2F) by ~2-fold (from 9% to ~18%) and expression of multiple pluripotency markers, including *DPPA2, DPPA4,* ZFP42 (*REX1*) and *NANOG* by ~2-6 fold (p <0.05) (Supplementary Fig. 3C,D), as compared to siControl, thus suggesting that AJSZ generally restricts HDFs ability to undergo cell fate reprogramming.

Given that AJSZ are broadly expressed in human adult tissues (https://www.proteinatlas.org/)^42^, we next asked whether their fate-stabilizing role would be conserved in non-fibroblast cell types, such as endothelial cells. First, we confirmed that AJSZ are expressed in primary human aortic endothelial cells (HAECs) (Supplementary Fig. 4A) and that their expression can be efficiently suppressed by specific siRNAs (Supplementary Fig. 4B). Finally, we evaluated the role of AJSZ in the CR assay in HAECs and found that AJSZ KD in combination with MGT overexpression significantly enhanced cardiac gene expression (2–10-fold, p<0.05) on day 3 (Supplementary Fig. 4C, D) and significantly increased the generation of ACTN1+ iCMs by ~1.5-fold (p<0.05) on day 20 (Fig. 2H, I) as compared siControl. In conclusion, our results collectively demonstrate that AJSZ play a lineage and cell type independent role in the promotion of cell fate stability in differentiated cells.

### AJSZ-mediated fate-stabilizing mechanisms are active at ground state in fibroblasts

To first gain insight into AJSZ mode of action, we determined whether the timing of AJSZ KD relative to reprogramming TFs overexpression would influence reprogramming efficiency (Supplementary Fig. 5A). Strikingly, while siAJSZ transfection on day −1 relative to MGT induction robustly enhanced CR efficiency, while AJSZ KD at day +1 failed to enhance CR efficiency (Supplementary Fig. 5B, C). Interestingly, AJSZ KD on day −2 or −3 before MGT overexpression progressively reduced the ability of AJSZ KD to improve CR efficiency (Supplementary Fig. 5D, E). Thus collectively, these results show that AJSZ-mediated fate stabilizing mechanisms are actively deployed at ground state in fibroblasts, prior to the TF-induced cell fate challenge.

### AJSZ bind to open and closed chromatin in a regionalized fashion at ground state in HDFs

To characterize the molecular nature of AJSZ-mediated fate-stabilizing mechanisms at ground state, we examined AJSZ interaction with the DNA in unchallenged HDFs using ChIP-seq (Fig. 3A). This approach identified 91,196 replicated peaks (binding sites) for JUNB; 44,100 for ATF7IP; 19,169 for SP7 and 4135 for ZNF207 across two duplicates (Fig. 3B and Supplementary Table 3), thereby revealing the pervasive association of the fate-stabilizing TFs with the chromatin at ground state, where JUNB and ATF7IP contributed to >85% of binding events. To further characterize these interactions, we next generated a genome-wide chromatin accessibility profile of HDFs at ground state using ATAC-seq and determined AJSZ ability to bind to open and closed chromatin. Remarkably, this analysis revealed that JUNB and ZNF207 preferentially bound to regions of open chromatin (78% and 94% of total binding respectively), while ATF7IP and SP7 mainly bound to closed chromatin (79% and 97% of total binding respectively) (Fig. 3C). In addition to these main interactions, 21% of ATF7IP (~9000 binding sites) and 22% of JUNB binding (~20,000 binding sites) occurred in open and closed chromatin respectively, thus revealing ATF7IP and JUNB ability to interact with both chromatin states albeit with differential frequency. In sum, our data show that at ground state, the four fate-stabilizing TFs interact with the DNA in a regionalized manner, with ATF7IP, JUNB and ZNF207 engaging the open chromatin and ATF7IP, JUNB and SP7 the closed chromatin.

**Fig. 3.**
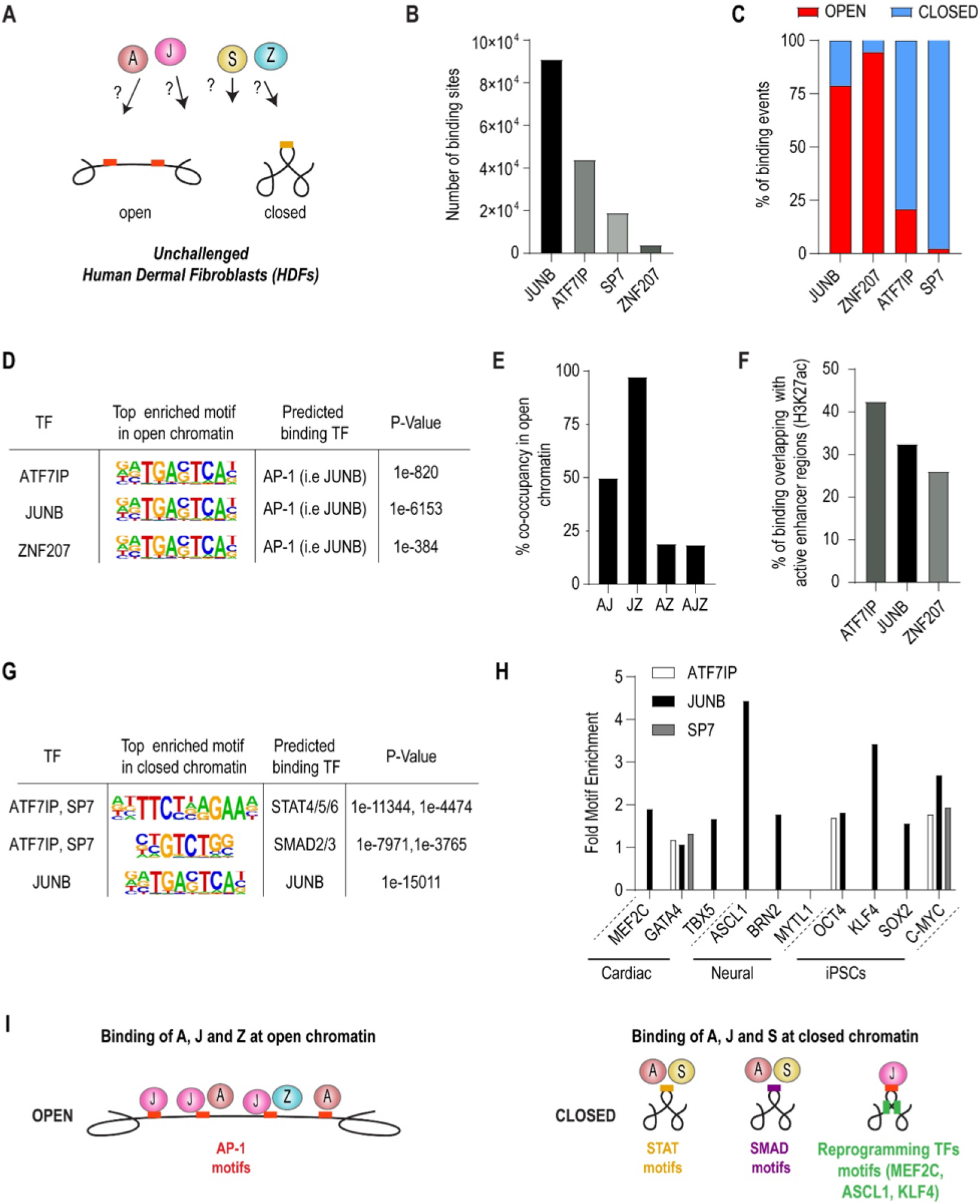
AJSZ binding properties in open and closed chromatin in HDFs. **(A)** Schematic showing that the binding of AJSZ in open and closed chromatin is unknown. ChIP-seq in human dermal fibroblasts (HDFs) using pulldowns of AJSZ was combined with scATAC-seq data in HDFs to identify where AJSZ bind. **(B)** Number of binding sites determined by ChIP-seq for each factor. Peaks were merged from 2 samples. **(C)** Percentage of binding sites located in open or closed chromatin. scATAC-seq was used to define closed chromatin. **(D)** Top enriched motif in the open chromatin binding sites for AJZ are the AP-1 motif. **(E)** Overlap in peak sites between combinations of AJZ show that JZ co-occupancy sites are almost all located in open chromatin. **(F)** AJZ individually show binding in active enhancer regions identified by the H3K27ac mark. **(G)** Top enriched motifs in closed chromatin indicate that A and S bind STAT and SMAD family motifs, while JUNB binds known JUNB motif. **(H)** Fold motif enrichment of AJS at LDFs motif sites indicates that AJS may block binding of these in ground state fibroblasts. **(I)** Schematic summarizing results showing AJZ bind AP-1 motifs in open chromatin while in the closed chromatin, ATF7IP and SP7 bind STAT and SMAD motifs with JUNB binding near key LDF (MEF2C, ASCL1, KLF4) motifs to possibly prevent expression of targets of these LDFs.

To delineate ATF7IP, JUNB and ZNF207 interactions at open chromatin regions, we performed a motif enrichment analysis and identified prototypical AP-1 TF recognition elements (-TGACTCA-)^43^, including putative JUNB binding motif, as top enriched sequences for all three barrier TFs (Fig. 3D and Supplementary Table 4). Consistent with these results, co-occupancy analysis revealed that 50% of ATF7IP and 97% of ZNF207 binding sites were co-occupied by JUNB (Fig. 3E and Supplementary Table 5), thus implying that JUNB might contribute to recruit ATF7IP and ZNF207 at these AP-1 motif-enriched chromatin regions. Next, to identify a potential role for ATF7IP, JUNB and ZNF207 interactions with the open chromatin, we asked whether these interactions would overlap with active enhancer regions (H3K27ac (ENCODE)). This analysis revealed that 26-42% of ATF7IP, JUNB and ZNF207 binding sites in the open chromatin overlapped with active enhancers marks (Fig. 3F), thus suggesting a role for these interactions in the regulation of transcription in HDFs at ground state. Finally, we noted that a significant portion (58-74%) of ATF7IP, JUNB and ZNF207 binding in open chromatin occurred outside of these active enhancer regions, thus suggesting a potential structural role for these interactions in the maintenance of these regions in an open state.

In regions of closed chromatin, ATF7IP and SP7 binding sites were most enriched for STAT4/5/6 and SMAD2/3 motifs, while JUNB binding sites were most enriched for AP-1 TFs motifs as in open chromatin (Fig. 3G and Supplementary Table 6). Co-occupancy analysis also revealed that 70% of SP7 binding sites were co-occupied by ATF7IP, while less than 2% of ATF7IP and SP7 binding sites were co-occupied by JUNB (Supplementary Table 7), thus indicating that ATF7IP and SP7 on the one hand and JUNB on the other, bind to distinct closed chromatin domains. To identify a potential role for these interactions, we next asked whether ATF7IP, JUNB and SP7-bound closed chromatin might be enriched for putative reprogramming TFs motifs. Remarkably, this analysis uncovered that JUNB-bound closed chromatin was significantly enriched for cardiac (MEF2C 1.89 fold, p-value=1e-172 and TBX5 1.66 fold, p-value=1e-366), neural (ASCL1 4.43 fold, p-value=1e854) and pluripotency (KLF4 3.42-fold, p-value=1e-71; cMYC 2.68 fold, p-value=1e-101) reprogramming TF motifs (Fig. 3H), thus suggesting that JUNB binding to AP-1 motifs might contribute to maintain these regions in a closed state, thereby limiting reprogramming TFs access to their target DNA. In summary, our results show that in unchallenged HDFs, AJSZ interact with both states of the chromatin in a genome-wide and regionalized fashion (Fig. 3I).

### AJSZ via JUNB binding to AP-1 motifs regulate chromatin accessibility dynamics during cell fate reprogramming

AJSZ extensively bind to both open and closed chromatin at ground state, thus we next asked whether these pre-existing interactions might contribute to limit chromatin remodeling induced by reprogramming TFs during the fate conversion process^44–47^. To address this question, we generated single-cell (sc) chromatin accessibility profiles from HDFs, 2 days after cardiac reprogramming TFs (MGT) overexpression in siControl- or siAJSZ-transfected HDFs using we scATAC-seq. t-distributed stochastic neighbor embedding (t-SNE) clustering of HDFs in MGT + siControl cells (15,859 cells), showed that cell clusters were distributed as a compact continuum (Fig. 4A), as in HDFs at ground state (Supplementary Fig. 6A), thus indicating that MGT overexpression alone did not induce major chromatin accessibility profile differences. In contrast, t-SNE clustering of MGT + siAJSZ cells (8966 cells) identified a discrete cell population (*cluster 2*) with a chromatin accessibility profile that significantly diverged from the remaining cell populations (black arrow, Fig. 4B). This cluster represented ~13% of total cells, which was notably similar to the percentage of ACTN1+ iCMs (~16%) generated after siAJSZ transfection and MGT overexpression in the CR assay (see Fig. 3C). Next, to define whether *cluster 2* cells were undergoing cell fate reprogramming, we performed an ontology analysis for genes with differentially accessible (DA) transcriptional start sites (TSSs) in *cluster 2* as compared to *clusters 1* and *3–7*, and observed a 5–10-fold enrichment (p<0.0001) for cardiac terms (*i.e.* striated muscle contraction and myofibril assembly) (Fig. 4C and Supplementary Table 8), involving a wide array of cardiac-specific genes (*i.e. NKX2-5, ACTA1,* and *NPPA*, Supplementary Fig. 6B). Collectively these results indicate that *1) cluster 2* represents a cell population with a chromatin accessibility profile indicative of cells undergoing fate reprogramming towards the cardiac lineage and *2)* that AJSZ regulate chromatin accessibility dynamics during this process.

**Fig. 4.**
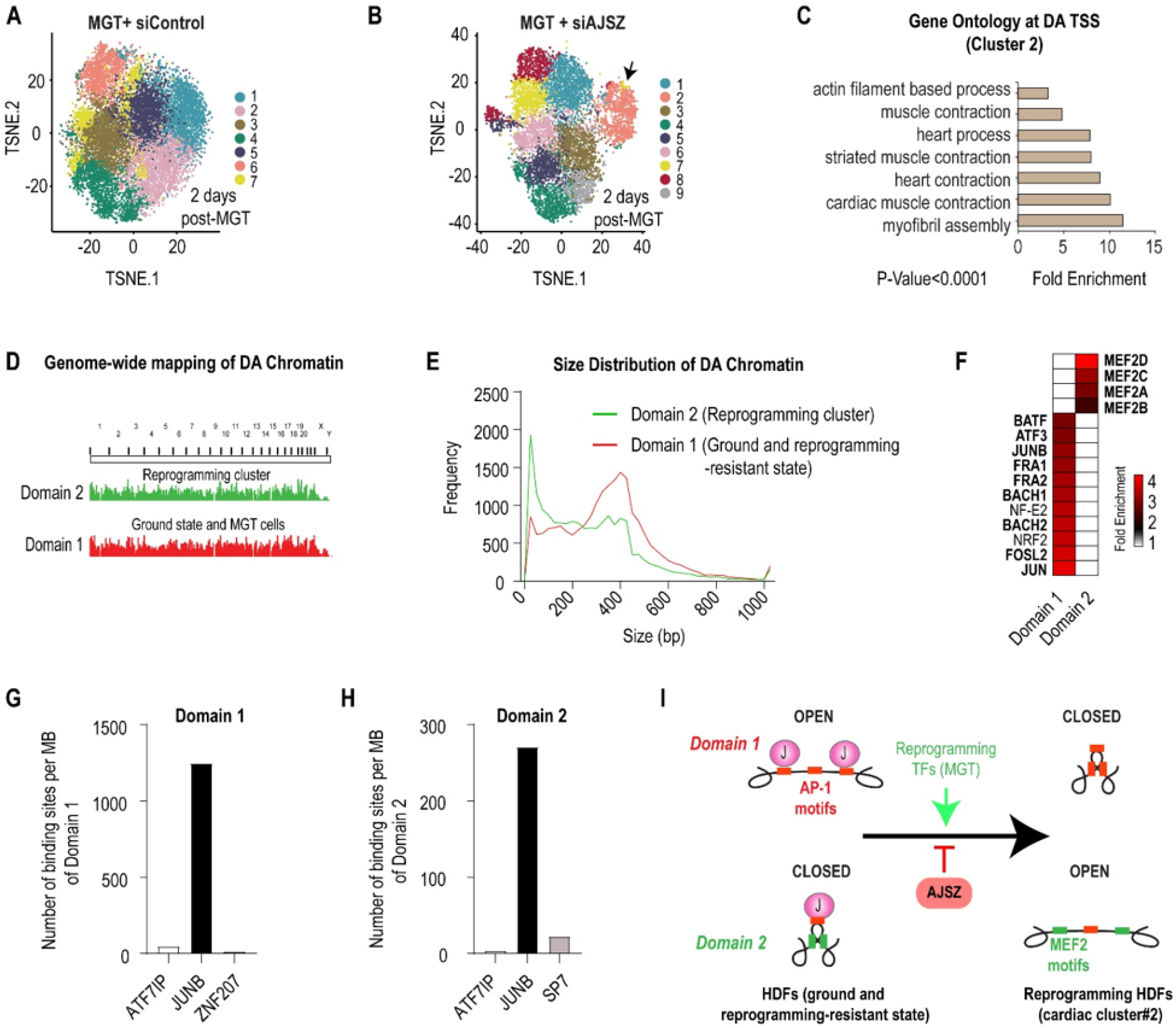
AJSZ epigenetically control expression of downstream barriers and agonists of cell fate reprogramming. **(A-B)** t-SNE visualization of cell clusters after scATAC-seq of siControl-transfected HDFs 2 days after mMGT overexpression (A), or siAJSZ-transfected HDFs 2 days after MGT overexpression (B). In contrast to the homogeneity of cells after MGT induction alone (A), knockdown of AJSZ prior to MGT overexpression (B) results in an open chromatin cardiac cluster (2, black arrow). **(C)** Gene Ontology analysis of genes with differential accessibility at the transcriptional start site (TSS) between Cluster 2 and all other clusters show enrichment of cardiac contraction related genes. **(D)** Density of *domain 1* and *domain 2* binding show that differential accessibility occurs throughout the genome. **(E)** Cluster 2 cells Distribution of *domain 1* and 2 region lengths indicating most regions are 200 to 500 bp long. **(F)** Comparison of motif enrichment between Domains 1 and 2. Motif analysis of *domain 2* indicates MEF2 family members bind. Motifs in *domain 1* are enriched for factors that bind AP-1 motif. **(G-H)** AJSZ binding from the ChIP-seq data in *domain 1* (G) and 2 (H). *domain 2* is highly enriched for AJSZ binding sites. **(I)** Model in which AJSZ via genome-wide JUNB binding to AP-1 motifs in *domain 1* chromatin impairs the ability of the reprogramming TF MEF2C to remodel chromatin and promote *domain 2* opening, thereby opposing cell fate conversion.

To further delineate the role of AJSZ in the regulation of DNA accessibility during cell fate reprogramming, we next mapped all regions of DA chromatin in cells with endogenous AJSZ levels in a non-reprogrammed or a reprogramming-resistant state (unchallenged HDFs and siControl+MGT HDFs; *domain 1*) as compared to cells with reduced AJSZ levels and undergoing fate reprogramming (*cluster 2* cells from siAJSZ+MGT HDFs; *domain 2*). Interestingly, this approach identified two domains of DA chromatin scattered across the entire genome (Fig. 4D), and consisting in short stretches of DNA (<600 bp) (Fig. 4E and Supplementary Fig. 6C), collectively spanning ~7 Mbp for *domain 1* and ~4.5 Mbp for *domain 2*. Next, to characterize potential TF motif composition differences between these two domains, we performed a differential enrichment analysis and observed a ~4-fold enrichment for AP-1 TFs motifs (*i.e.* JUNB putative binding site, 1 motif/kb), in *domain 1* as compared to *domain 2* (Fig. 4F, Supplementary Fig. 6D and Supplementary Table 9). In contrast, in *domain 2*, MEF2 motifs were the most differentially enriched (~3.5-fold, 0.6 motif/kb) (Fig. 4F) and included the canonical binding sequence^48^ for CR TF: MEF2C (Supplementary Fig. 6E and Supplementary Table 9).

Finally, to determine whether the fate-stabilizing TFs might directly bind to these domains, we quantified the number of AJSZ binding sites at *domain 1* and 2 at ground state in HDFs. This analysis revealed that JUNB was the major interactor for both domains and was involved in >88% of binding events (Fig. 4G,H), thus suggesting that JUNB plays a direct role in the establishment or maintenance of *domain 1* in an open state and *domain 2* in a closed state. Collectively, our data show that AJSZ, *via* direct JUNB binding to AP-1 motif-enriched chromatin, contribute to limit the number of motifs accessible to the reprogramming TFs, thereby restricting their ability to bind to their target DNA and promote cell fate conversion (Fig. 4I).

### AJSZ play a genome-wide and proximal role in the regulation of transcription during cell fate reprogramming

TFs are proximal regulators of transcription28,49 and consistent with this role, our ChIP-Seq data also revealed that AJSZ bind to > 9300 promoter-TSS regions (−1 kbp, +0.1 kbp) in HDFs at ground state (Supplementary Table 10). The magnitude of this interaction led us to postulate that, in addition to their role in the regulation of chromatin accessibility, AJSZ might also oppose cell fate reprogramming via the proximal regulation of transcription. To test this hypothesis, we performed genome-wide RNA-seq of control and AJSZ KD HDFs 2 days after cardiac reprogramming TFs (MGT) overexpression. We identified 736 differentially expressed (DE) genes, of which 501 and 235 were downregulated and upregulated, respectively by AJSZ KD (p<0.05, Supplementary Table 11 and Fig. 5A). Integration of RNA- and ChIP-seq datasets revealed that ~2/3 of the DE genes (460 of 736, Supplementary Table 12) were bound at their core promoter regions by at least one of the 4 fate-stabilizing TFs in HDFs (Fig. 5B). Notably, core promoter binding correlated with gene downregulation for ~75% of the DE genes, thus indicating a predominantly activating role for AJSZ in the regulation of transcription during cell fate reprogramming. In this context, GO analysis of core promoter-bound DE genes revealed a significant enrichment of terms related to cell fate specification, cardiac muscle differentiation, fibroblast proliferation, collagen organization and TGF-βsignaling, substantiating the potential involvement of AJSZ in the control of cell fate-regulating transcriptional programs in fibroblasts (Fig. 5C and Supplementary Table 13). Moreover, assessment of individual AJSZ contributions to core promoter binding revealed that 97% of the core promoters were bound by JUNB in HDFs (Fig. 5D). In this context, analysis of JUNB binding site distribution showed that these interactions were centered at the TSS regions (Fig. 5E) and could be observed for both downregulated (*TAGLN*) and upregulated (*MEF2C*) genes (Fig. 5F, G). Collectively, these results show that AJSZ play a proximal role in the regulation of fibroblast (TGF-β, collagen organization, proliferation) and cell fate-modulating transcriptional programs during reprogramming, *via* JUNB binding to TSS regions.

**Fig. 5.**
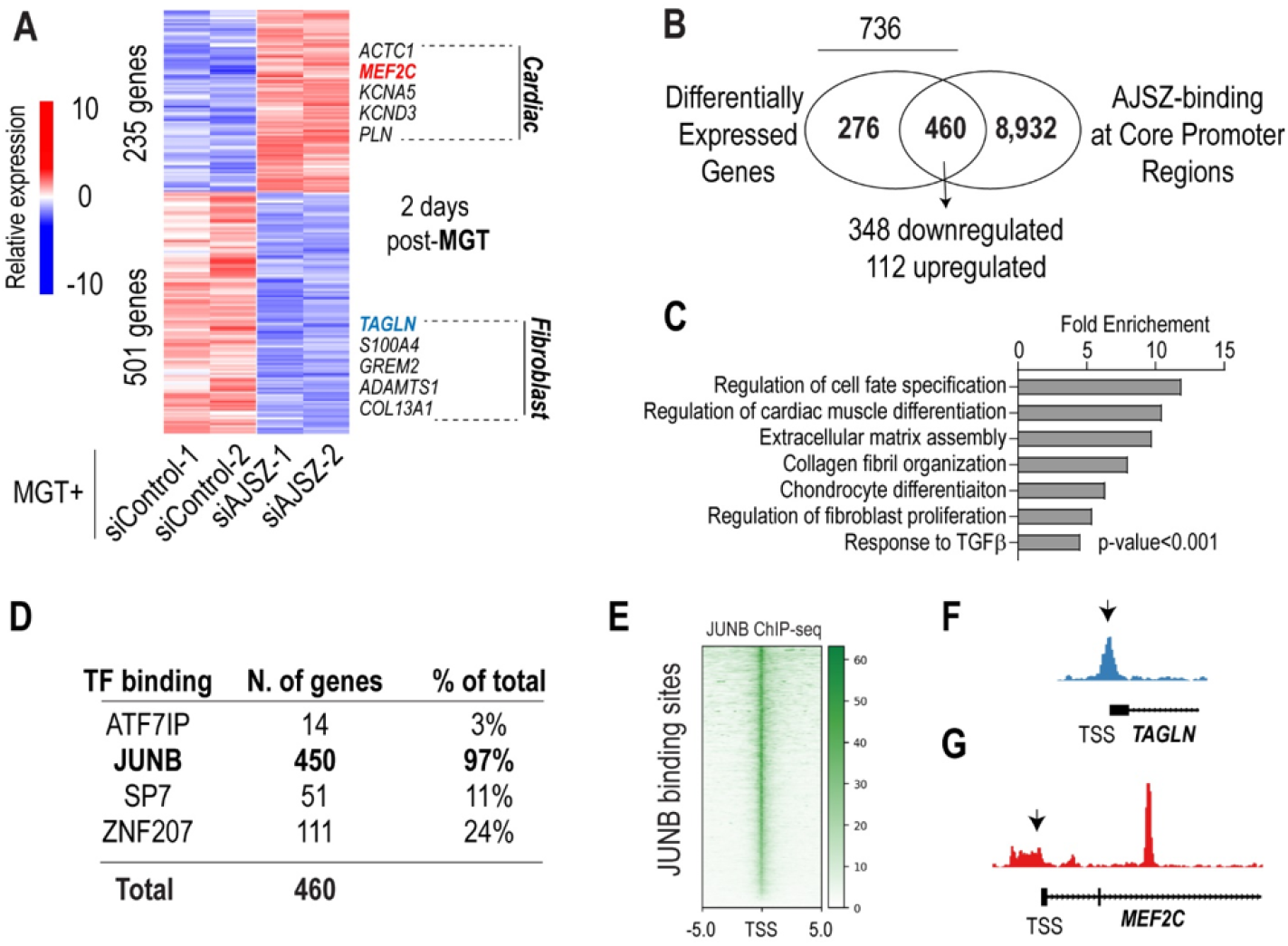
AJSZ differentially regulate gene expression cell fate reprogramming. **(A)** Heatmap of differentially expressed (DE) genes in siControl- and siAJSZ-transfected HDFs 2 days after MGT overexpression. **(B)** Venn diagram showing overlap between DE and core promoter bound (−1 kb-TSS-+0.1 kb) genes. 460 genes were both DE and bound by AJSZ at core promoter regions, including 348 downregulated and 112 upregulated genes. **(C)** Bar charts showing top ranked biological process terms enriched for the 460 DE and core promoter bound genes. **(D)** Breakdown of the percentage of DE and core promoter bound genes containing A, J, S, and/or Z binding sites. **(E)** ChIP-seq track for JUNB at JUNB binding sites. **(F, G)** Genome browser views showing binding of JUNB at TAGLN (f) and at MEF2C core promoter regions (G) in HDFs.

### AJSZ promote cell fate stability via the transcriptional repression of a conserved network of genes required for the reprogramming process

Given that AJSZ act both as fate stabilizers and proximal regulators of transcription during cell fate reprogramming, we next postulated that they might promote cell fate stability by upregulating reprogramming barrier genes (Fig. 6A), while downregulating genes required for reprogramming (= agonists) (Fig. 6E). To investigate this, we tested top 25 percentile core promoter-bound and downregulated genes, in MGT+siAJSZ condition as compared to MGT+siControl (Supplementary Table 14), for barrier to reprogramming function using a siRNA-mediated KD strategy in the iMGT-MEF CR assay. Consistent with our hypothesis, this approach identified two hits, siChst2 and siNceh1, that robustly enhanced CR efficiency (1.5- and 2-fold, p<0.05 and p<0.01, respectively; Fig. 6B–D), thus uncovering a novel barrier to CR roles for carbohydrate sulfotransferase 2 (*Chst2*) which mediates 6-O sulfation within proteoglycans^50^, and neutral cholesterol ester hydrolase 1 (*Nceh1*) which regulates lipid droplet formation^51^ and platelet-activating factor synthesis^52^.

**Fig. 6.**
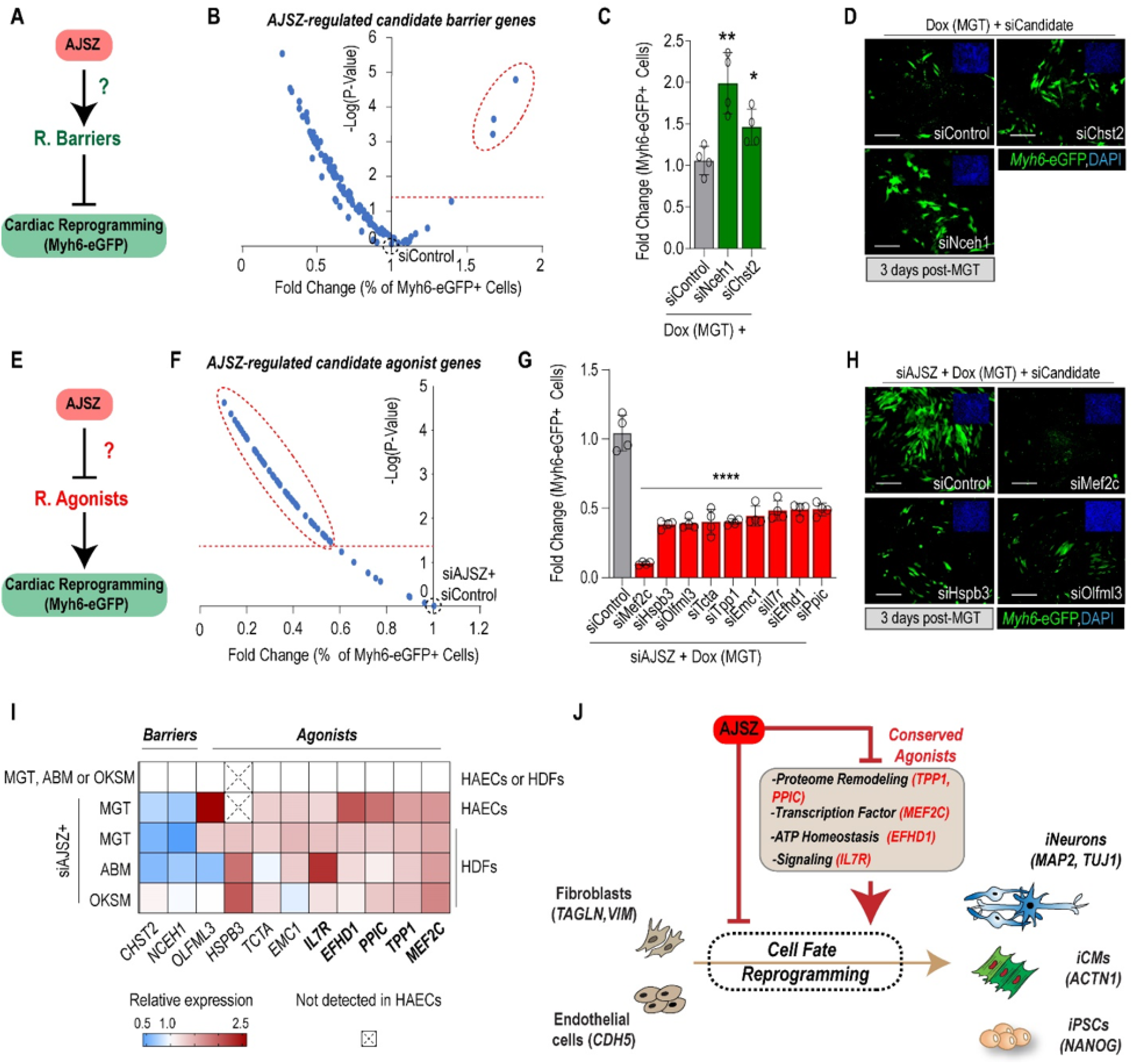
AJSZ epigenetically control expression of downstream barriers and agonists of cell fate reprogramming. **(A)** Schematic showing the hypothesis that AJSZ positively regulate expression of reprogramming barriers. **(B)** Volcano plot showing the screening results for siRNAs directed against top 25 percentile downregulated (MGT+siAJSZ vs MGT) and core promoter bound genes in the iMGT-MEF CR assay. The top 3 candidate barrier genes are circled. N=4 per condition. **(C)** Histogram showing validation of siNceh1 and siChst2 effect on CR. **(D)** Representative images for siControl, siChst2 and siNceh1 conditions. Myh6-eGFP+ cells are shown in green and cell nuclei are stained blue (DAPI, top right insets). **(E)** Schematic of the hypothesis that AJSZ negatively regulate reprogramming agonists in HDFs. **(F)** Volcano plot showing the screening results for siRNAs directed against top 77 upregulated (MGT+siAJSZ vs MGT) and core promoter bound genes in siAJSZ-induced iMGT-MEF CR assay. **(G)** Histogram showing validation of top 9 siRNAs that blunt siAJSZ-induced CR without affect cell viability. (H) Representative images for siAJSZ+ siControl, siMef2c, siHspb3, or siOlfml3 conditions. **(I)** Heatmap summarizing AJSZ expression dependence of identified barriers and agonists in HAECs and HDFs, 2 days after MGT, ABM or OKSM overexpression. **(J)** Cell fate and cell type-independent model for the transcriptional regulation of cell fate reprogramming by AJSZ. Scale bars: 50 μm. Student’s t-test. *p<0.05, **p<0.01, ****p<0.0001.

Next, to identify AJSZ-regulated reprogramming agonists (Fig. 6E), we tested siRNAs against top 25 percentile core promoter-bound and upregulated genes, in MGT+siAJSZ condition as compared to MGT+siControl (Supplementary Table 15), for reprogramming requirement in the siAJSZ-induced CR assay. This approach identified 61 siRNAs that reduced CR efficiency induced by AJSZ KD by 50% (Fig. 6F); of these, only 9 siRNA hits had no effect on cell viability and were selected for further analysis (Fig. 6G, H). Notably, the 9 agonists could be grouped into 4 gene categories involved in regulation of cell fate specification (Mef2c)^48^, protein folding and degradation (Emc1, Hspb3, Ppic, Tpp1)^53–55^, inflammation and TGF-βsignaling (Il7r, Olfml3, Tcta)^56–58^, and ATP homeostasis (Efhd1)^59^.

Finally, we asked whether the AJSZ-mediated regulation of reprogramming agonists and barriers identified above, was lineage and/or cell type-specific. Remarkably, consistent with a conserved function of AJSZ as fate stabilizers, AJSZ KD led to a lineage- and cell type-independent upregulation of reprogramming agonists (*MEF2C, TPP1, PPIC, IL7R,* and *EFHD1*) after 3 days of reprogramming (Fig. 6I). Collectively, these results show that a conserved component of the AJSZ-mediated regulation of cell fate stability is to proximally repress genes involved in the promotion of large-scale phenotypic changes (*i.e.* proteome remodeling, cell fate specification, ATP homeostasis, inflammation signaling) (Fig. 6J).

### AJSZ KD Enhances MGT ability to Improve Cardiac Function and Reduce Fibrosis after MI

Retrovirus-mediated overex-pression of MGT has been shown to induce CR in fibroblasts *in vivo* and improve adult heart function post-myocardial infarction (MI)60,61. To assess AJSZ function *in vivo*, we first established that co-infection of fibroblasts with individual shRNAs targeting AJSZ could efficiently enhance CR *in vitro* (Supp Fig.7A). Next, to assess whether shRNA AJSZ might enhance MGT ability to improve cardiac function after MI, we permanently ligated the left anterior descending coronary artery in 10- to 12-week old mice and after 3 days, we delivered virus mixtures containing either MGT or MGT+shRNAs at the site of injury post -MI, using ultrasound-guided injection (Fig. 7A and Supp Movie s.2)^62^. We next assessed cardiac function in the PBS, polycistronic MGT-, and MGT+shAJSZ-injected mice by echocardiography at 2 and 4 weeks after MI in a blinded fashion. Remarkably, ejection fraction (EF) and fractional shortening (FS) of MGT+shAJSZ-injected left ventricle was increased by 250% as compared to PBS-injected and 50% as compared MGT-injected mice at 2 and 4 weeks after MI (Fig. 7B-D and Supp Fig. 7C,D). We, next conducted histological analyses to quantify the scar size after 4 weeks by Masson trichrome staining. Quantification on serial sections, revealed a ~40% reduction in scar area in MGT + shAJSZ. as compared to MGT (Fig. 7E,F). Collectively, these results demonstrate that KD of AJSZ is sufficient to improve heart function as compared to MGT alone and also demonstrate that the targeted inactivation of cell fate stabilizers represents a viable strategy to improve adult function post-injury.

**Fig. 7.**
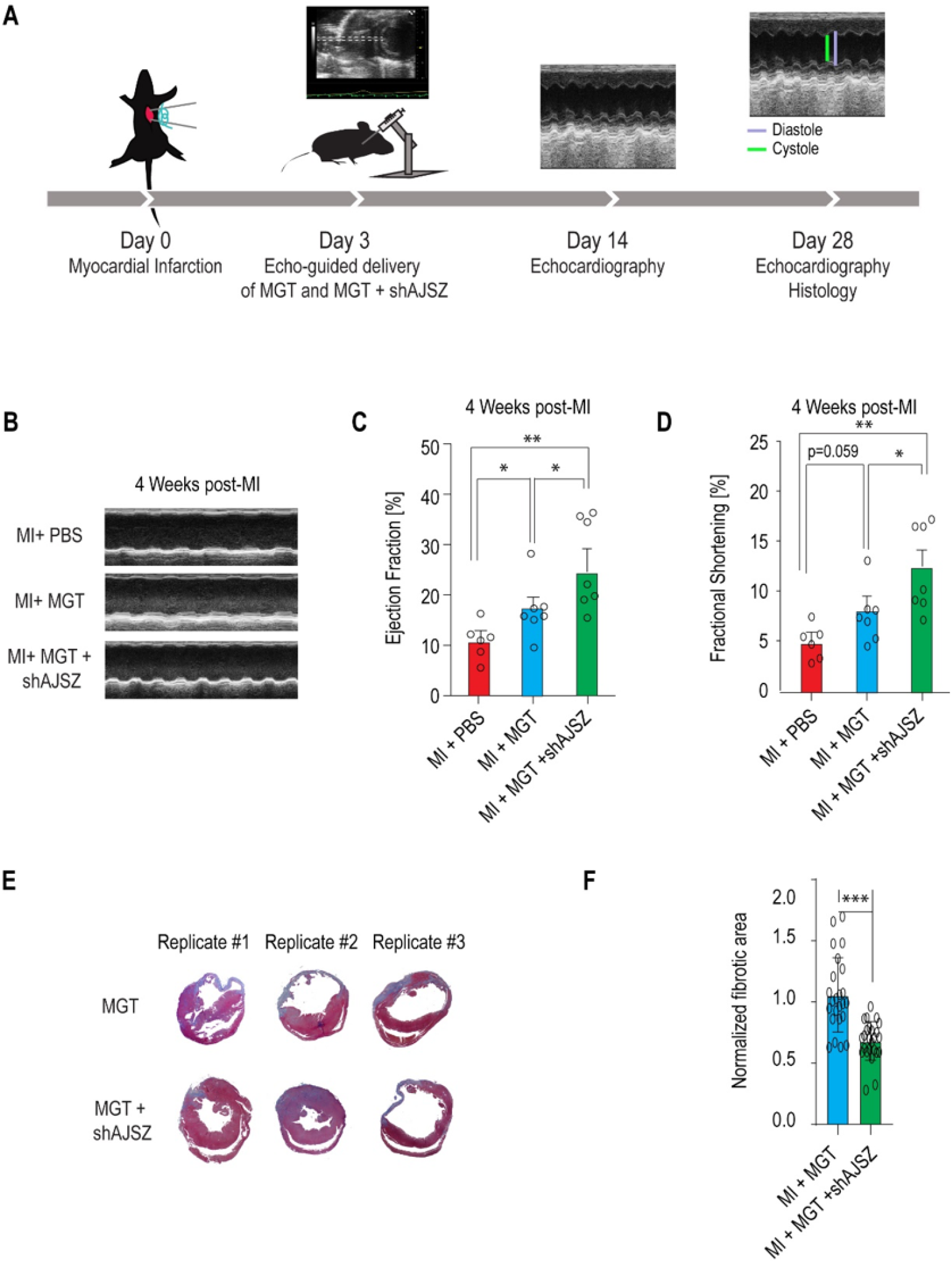
AJSZ KD in combination with MGT overexpression improves heart function post-MI. **(A)** Schematic depicting experimental strategy to test the role of AJSZ on heart function post-MI. **(B)** Examples of echocardiograms from MI+PBS, MI+MGT or MI+MGT+shAJSZ used to measure ventricular contractility 4 weeks after MI. **(C)** Ejection fraction (EF) and **(D)** fractional shortening (FS) of the left ventricle were serially quantified by echocardiography in mice injected with PBS, MGT and MGT+ shAJSZ (n = 7 for each group). Cardiac function was improved with MGT+ shAJSZ as compared to PBS or MGT conditions. **(E)** Representative histological heart sections of three different mice for MGT or MGT+shAJSZ condition, with Masson trichrome staining and quantification **(F)** showing reduced fibrosis area in MGT+shAJSZ as compared to MGT mice. Groups were compared using two-tailed unpaired. *P < 0.05, **P < 0.01 and ***P < 0.001.

## Discussion

Here, we report on the identification of four TFs (AJSZ), which collectively promote cell fate stability in terminally differentiated cells and oppose TF-mediated fate reprogramming in a lineage- and cell type-independent manner. A detailed analysis of AJSZ mode of action revealed that they limit cell fate reprogramming by concomitantly (i) modulating chromatin accessibility and impairing reprogramming TFs to access their target DNA in a genome-wide fashion; and (ii) by restricting the expression of genes required for large-scale phenotypic changes. Remarkably, targeting these 4 fate stabilizing TFs *in vivo* was sufficient to significantly enhance MGT-induced heart repair and thus, identifying fate stabilizers as a new promising class of targets to enhance adult organ regeneration.

### Identification of a generic mechanism regulating cell fate stability in differentiated cells

Given the diversity of cell types co-existing in multicellular organisms and the necessity for these cells to maintain a differentiated phenotype to fulfill specialized functions, a central point to discuss here, is whether AJSZ’s fate-stabilizing role is generic or specific to a subset of differentiated cells and/or lineages. Strikingly, results from our multi-model systems approach (Figure 1 and 2) show that the role of AJSZ as barriers to cell fate reprogramming is conserved across multiple species (mouse and human), cell types (fibroblasts and endothelial cells) and lineages (cardiac, neural, iPSCs). Moreover and consistent with a general fate-stabilizing role, AJSZ are expressed in most adult tissues (https://www.proteinatlas.org/)42, while their expression is reduced^63–67^ in fate-destabilized cancer cells from distinct lineages (*i.e.* blood, bone, prostate). Finally, expression profiling of AJSZ in hiPSCs reveals that JUNB and SP7 are not expressed in these undifferentiated cells (Supplementary Fig. 8A), while their differentiated progeny robustly expresses all four factors in their nuclei (Supplementary Fig. 8b). Thus, given their 1) conserved role as barriers to cell fate reprogramming, 2) elevated and ubiquitous expression in differentiated cells and 3) diminished expression in fate-destabilized cancer and undifferentiated cells, we propose that AJSZ may contribute to establish a generic mechanism mediating the promotion of cell fate stability in differentiated cells. In this context, we hypothesize that yet to be identified regulators of AJSZ expression and/or activity, might represent novel factors mediating phenotypic stability in differentiated cells.

### Regionalized DNA binding mediates integrated control of chromatin accessibility and transcription during cell fate reprogramming

TFs recognize specific DNA sequences to control chromatin architecture46 and transcription28, and here we explored how AJSZ integrate these two regulatory dimensions to mediate cell fate stability. Our detailed analysis (ChIP-, scATAC- and RNA-seq and ENCODE, Figure 3, 4 and 5) of AJSZ mode of action at ground state and during cell fate reprogramming, revealed that AJSZ exert their role via direct DNA binding to 3 distinct chromatin regions. - The first region involves A, J and Z binding to AP-1 motif-enriched open chromatin, where they fulfill two distinct roles: 1) to proximally regulate the expression of lineage-appropriate (*i.e.* TGF-β, collagen organization, proliferation in HDFs) and barrier network genes and 2) to maintain a subset of A, J and Z-bound chromatin (*domain 1*) in an open state. Although, more work will be needed to understand how A, J and Z binding to *domain 1* contributes to maintain these regions in an open state and oppose cell fate reprogramming, recent findings from Li et al 44 have shown that appropriate closing of AP-1 motif-enriched chromatin is required for efficient reprogramming of MEFs into iPSCs. - The second main region of interaction involves JUNB binding to reprogramming TF motifs-enriched (*i.e.* MEF2C, ASCL1, KLF4, cMYC) closed chromatin (Fig.3). In this context, our comparative chromatin accessibility analysis (siControl vs siAJSZ) during cardiac reprogramming of HDFs (Fig.4), shows that under control conditions, pre-existing binding of JUNB to cardiac reprogramming TF (MEF2C) motif-enriched regions (*domain 2*), remains in a closed state even when MGT is overexpressed, thus contributing to limit MEF2C access to target DNA and thereby opposing cell fate reprogramming. Conversely, reduced AJSZ levels enable the de novo opening of chromatin from *domain 2* and the generation of cells with a cardiac-like chromatin accessibility profile. In this context, it remains to be established how JUNB binding to *domain 2* contribute to maintain these regions in a closed state. - Finally, the third region of interaction involves ATF7IP and SP7 binding to closed chromatin regions enriched for SMAD and STAT motifs. Interestingly, previous studies have shown that TGF-β/SMAD inhibition enhances iPSCs^68^ and cardiac^22,47^ reprogramming efficiency from fibroblasts and while STAT signaling impairment increases Ascl1-induced transdifferentiation of glial cells into neurons and regeneration in adult mouse retina^69^. In this context, we propose that ATF7IP and SP7 binding at SMAD and STAT motif-enriched closed chromatin contribute to mediate TGF-β/SMAD and STAT signaling and thereby promoting cell fate stability. In sum, motif-specific binding of AJSZ to 3 distinct chromatin regions enables to regulate chromatin accessibility and transcription on a genome-wide scale to promote cell fate stability in differentiated cells.

### AJSZ transcriptionally control genes regulating cells ability to undergo large-scale phenotypic changes

The delineation of genes and pathways regulating cell fate stability and cells ability to undergo large-scale phenotypic changes is essential for our ability to understand how cells normally resist cell fate challenges (*i.e.* cancer, virus infection), but also can be used as a diagnostic markers to monitor cells undergoing oncogenic transformation or conversely as a molecular tool to promote engineered reprogramming. Here, in addition to chromatin accessibility regulation, the systematic functional evaluation of AJSZ-regulated genes during the process of cell fate conversion (Fig.6J), revealed the existence of a conserved network of agonists that limit cells ability to undergo reprogramming. A remarkable property for these genes, is their functional requirement for the reprogramming process downstream of AJSZ KD, thus implying that they act as a non-redundant fate-stabilizing mechanism in addition to the AJSZ-mediated regulation of chromatin accessibility described above. At the molecular level, these conserved agonists fall into four cellular ontologies: regulators of protein folding and degradation (*PPIC, TPP1*), cell fate specification (*MEF2C*), ATP generation (*EFHD1*) and inflammation signaling (*IL7R*), thus, suggesting that these molecular functions represent rate-limiting steps restricting cells ability to undergo large-scale phenotypic changes and thus fate reprogramming. Finally, and consistent with a role for these genes in the promotion of phenotypic destabilization, upregulation and/or gain-of-function mutations for all 5 genes is found in cancer cells originating from all three germ layers ^57,70–75^.

### Targeting AJSZ, a promising new technology for regenerative medicine

The use of direct cell fate reprogramming to promote adult organ repair is a promise of regenerative medicine13, however, to date low reprogramming efficiency still limits application to the clinic12. In this study, we aimed at evaluating whether inhibition of fate-stabilizing mechanisms might enhance reprogramming TFs ability to improve adult organ function post-injury. Remarkably, our data (Figure 7) show that RNAi-mediated loss of AJSZ function in combination with MGT overexpression is sufficient to improve heart function post-MI by 50% as compared to treatment with MGT alone and 250% as compared to no treatment. In this context, this effect could be observed as early as 2 weeks post-treatment and was maintained for at least a month. Collectively, these results suggest that the inhibition of fate stabilizers activity might represent a novel target space to rapidly improve adult heart function post-injury. Although more work will be needed to further delineate the molecular mechanisms underlying *in vivo* regeneration, to our knowledge, this is the first study demonstrating a viable strategy to treat already infarcted hearts. In this context, it should also be tested whether AJSZ loss of function in combination with overexpression of other lineage-specific (*i.e.* neural) reprogramming TFs might enhance repair in other organs.

## Methods

### iMGT-Myh6-eGFP-MEFs and screening assays

Immortalized Dox-inducible iMGT-Myh6-eGFP-MEFs were described previously^35^. The cells were cultured in plates precoated with 0.1% gelatin (Stem Cell Technologies, 7903) and maintained in Fibroblast Culture Medium (FCM) consisting of DMEM (Corning, 10-013-CV), 10% fetal bovine serum (FBS; VWR, 89510-186), and 1% penicillin/streptomycin solution ((10,000 U/mL), Catalog : 15140122) at 37°C in a 5% CO2 atmosphere. One day prior to siRNA transfection (day - 1), cells were detached by addition of 0.25% trypsin-EDTA (Thermo Fisher, 25200056) for 3 min at 37°C, and then washed in FCM, centrifuged, and resuspended in Induced-CM Reprogramming Medium (iCRM), consisting of DMEM, 20% Medium 199 (Gibco, 11150-059), 10% FBS, and 1% P/S. Cells were plated in 384-well plates at 10^3^ cells/well and transfected with an siRNA library directed against 1435 mouse TFs (Dharmacon-Horizon Discovery; siGenome-siRNA library, G-015800). The next day (day 0), 1 μg/mL doxycycline hydrochloride (Dox; Sigma, D3072) diluted in iCRM was added to the cells to induce MGT expression. On day 3, the cells were fixed with 4% paraformaldehyde (PFA) and processed for immunostaining, microscopy, and Myh6-eGFP quantification. The top 20 siRNA hits were validated using independent siRNAs (Dharmacon-Horizon Discovery; ON-Target plus pooled siRNAs). All possible combinations of the top 8 siRNAs (255 combinations) were assembled by echo-spotting using an Echo 550 liquid handler (Labcyte) and the cells were processed as described above. All experiments were performed in quadruplicate. For follow-up analyses, the cells were collected on 3 after Dox addition for qRT-PCR analysis.

### Human primary dermal fibroblasts (HDFs)

Newborn human primary foreskin fibroblasts were obtained from American Type Culture Collection (ATCC; CRL-2097, CCD-1079Sk) and cultured in plates precoated with 0.1% gelatin in FCM. For cardiac reprogramming, the cells were allowed to reach 80% confluency, harvested using trypsin-EDTA as described above, resuspended in iCRM, added to 384-well plates at 10^3^ cells/well, and transfected with the indicated siRNAs. The next day (day 0), the cells were transduced by addition of 1 μL/well of mouse MGT retrovirus35 diluted in iCRM. Cells were collected on day 2 for RNA-seq and scATAC-seq experiments, on day 3 days for qRT-PCR experiments, and on day 30 for calcium handling assays. Immunostaining was performed on the days indicated in the legends. During the incubations, 50% of the iCRM medium was exchanged every other day up to day 8, and was then replaced with RPMI 1640 (Life Technologies, 11875093), 1% B27 supplement (Life Technologies, 17504044), and 1% P/S. Neuronal reprogramming was induced as previously described7,38. In brief, HDFs were cultured and harvested as described above, resuspended in FCM, added to 384-well plates at 3×10^3^ cells/well, and transfected with siRNAs. The next day (day −1), 0.25 μL of F-ABM lentiviral mix (1:1:1:1 of Addgene plasmids 27150, Tet-O-FUW-Ascl1; 27151, Tet-O-FUW-Brn2; 27152, Tet-O-FUW-Myt1l; 20342, FUW-M2rtTA) was added to each well, and the cells were incubated for an additional 24 h. The following day (day 0), ABM expression was induced by addition of 2 μg/mL Dox diluted in FCM. On day 2, media was replaced by Minimal Neuronal Medium consisting of DMEM/F-12 (Gibco, 11220-032), 1% B27, 1% N2 Supplement (Gibco, 17502-048), and 1% human recombinant insulin and zinc solution (Gibco, 12585-014). On day 3, cells were harvested and processed for immunostaining or qRT-PCR analysis. Induced pluripotent stem cell reprogramming was induced as previously described41. In brief, HDFs were cultured and harvested as described above, resuspended in FCM, added to 384-well plates at 1×10^3^ cells/well, and transfected with siRNAs. Cells settled to the bottom of the well for 5 minutes at room temperature and then 0.25 μL of F-OKMS lentiviral mix (1:1 of Addgene plasmids 51543, FUW-tetO-hOKMS; 20342, FUW-M2rtTA) was added to each well, and the cells were incubated for 24 h. The following day (day 0), OKMS expression was induced by addition of 2 μg/mL Dox diluted in FCM. On day 2, 4, and 6, media was replaced with Stem Cell Technologies mTeSR Plus Kit (Stem Cell Technologies 100-0276). On day 7, cells were harvested and processed for immunostaining or qRT-PCR analysis of pluripotency genes.

### Human adult aortic endothelial primary cells (HAECs)

Human adult aortic endothelial cells (HAECs) were obtained from ATCC (PCS-100-011) and cultured in plates precoated with 0.1% gelatin in Vascular Cell Basal Medium (VCBM; ATCC, PCS-100-030) containing endothelial cell growth kit-BBE (ATCC, PCS-100-040) and 0.1% P/S. When the cells reached 80% confluency, they were harvested with trypsin-EDTA solution for primary cells (ATCC, PCS-999-003) for 3 min at 37°C, washed with trypsin-neutralizing solution (ATCC, PCS-999-004), resuspended in VCBM/kit-BBE medium, added to 384-well plates at 3×103 cells/well, and transfected with siRNAs. One day later (day 0), cells were transduced by addition of 1 μL/well of mouse MGT retrovirus. The cells were collected on day 3 for qRT-PCR analysis and on day 20 for immunostaining. During the incubation, 50% of VCBM/kit-BBE medium was exchanged every other day starting on day 4.

### siRNA transfection

Mouse and human siRNAs were purchased from Dharmacon or Ambion and added at a final concentration of 25 nM. A negative control siRNA (referred to as siCTR or siControl) was obtained from Dharmacon. Transfection was performed using Opti-MEM (Gibco, 31985070) and Lipofectamine RNAiMAX (Gibco, 13778150) according to the manufacturer’s instructions. siRNA transfection was performed on day −1, unless otherwise noted.

### Immunostaining and fluorescence microscopy

Cells were fixed with 4% PFA in phosphate-buffered saline (PBS) for 30 min and blocked with blocking buffer consisting of 10% horse serum (Life Technologies, 26050088), 0.1% gelatin, and 0.5% Triton X-100 (Fisher Scientific, MP04807426) in PBS for 30 min. Cells were incubated with primary antibodies overnight at 4C, and followed by secondary antibodies with 4,6-diamidino-2-phenylindole (DAPI; Sigma, D9542) for 1 h in the dark at room temperature. Cells were washed with PBS between each step. Cell were imaged with an Image Xpress confocal microscope (Molecular Devices) and fluorescence was quantified with Molecular Device software. Experiments were performed in quadruplicate.

The primary antibodies used were: rabbit anti-ATF7IP (Sigma, HPA023505, 1:200); rabbit anti-ATF7IP (Invitrogen, PA5-54811, 1:200); rabbit anti-JUNB (Abcam, Ab128878, 1:200); rabbit anti-SP7/OSTERIX (Abcam, Ab22552, 1:500); mouse anti-ZNF207 (Sigma, SAB1412396, 1:500); rabbit anti-TAGLN (Abcam, Ab14106, 1:800); guinea pig polyclonal anti-Vimentin (Progen, GP53, 1:100); mouse anti-VIMENTIN (Santa Cruz Biotechnology, sc-373717, 1:800); goat polyclonal anti-TAGLN (GeneTex, GTX89789, 1:800); rabbit polyclonal anti-TNNT2 (Sigma, HPA017888, 1:100); mouse anti-ACTN1 (Sigma, A7811, 1:800); goat anti-PECAM1 (H3) (Santa Cruz Biotechnology, Sc1506, 1:200); rabbit anti-MAP2 (Abcam, Ab32454, 1:200); and mouse anti-TUJ1 (R&D Systems, MAB1195, 1:200); rabbit anti-NANOG (Abcam, ab109250, 1:200). Secondary antibodies were: Alexa Fluor 488 goat-anti-rabbit IgG (H+L) (Invitrogen, A11008, 1:1000); Alexa Fluor 488 donkey-anti-mouse IgG (H+L) (Invitrogen, A21202 1:1000); Alexa Fluor 488 goat anti-Guinea Pig IgG (H+L) (Invitrogen, A 11073 1:100); Alexa Fluor 568 goat-anti-mouse IgG (H+L) (Invitrogen, A10037, 1:1000); Alexa Fluor 568 donkey anti-goat IgG (H+L) (Invitrogen, A11057, 1:1000); Alexa Fluor 680 donkey-anti-mouse IgG (H+L) (Invitrogen, A10038, 1:1000); and Alexa Fluor 680 donkey-anti-rabbit IgG (H+L) (Invitrogen, A10043, 1:1000).

### RNA extraction and qRT-PCR

Total RNA was extracted using Zymo Research Quick-RNA MircoPrep Kit (Zymo Research, R1051) or TRIzol reagent (Invitrogen, 15596026) and chloroform (Fisher Chemical, C298-500) following the manufacturers’ recommendations. RNA in the aqueous phase was precipitated with isopropanol, centrifuged, washed with 70% ethanol, and eluted in DNase- and RNase-free water. RNA concentration was measured by Nanodrop (Thermo Scientific). Aliquots of 1 μg of RNA were reverse transcribed using a QuantiTect Reverse Transcription kit (Qiagen, 205314), and qPCR was performed with iTaq SYBR Green (Life Technologies) using a 7900HT Fast Real-Time PCR system (Applied Biosystem). Gene expression was normalized to that of glyceraldehyde 3-phosphate dehydrogenase (*GAPDH*) for human samples or β-actin (*Actb*) for mouse samples using the 2-,6.,6.Ct method. Human and mouse primer sequences for qRT-PCR were obtained from Harvard Primer Bank. Primers were: ATF7IP (#38261961c1), Atf7ip (#34328232a1), JUNB (#44921611c1), Junb (#6680512a1), SP7 (#22902135c2), Sp7 (#18485518a1), ZNF207 (#148612834c1), Zfp207 (#7212794a1), ACTC1 (#113722123c1), MYL7 (#50593014c1), NPPA (#23510319a1), NPPB (#83700236c1), RYR2 (#112799846c1), SCN5A (#237512981c1), TNNI3 (#151101269c1), TNNT2 (#48255880c1), GAD67 (#58331245c2), PVALB (#55925656c2), SYN1 (#91984783c1), vGLUT2 (#215820654c2), MYH6 (#289803014c3), HSPB3 (#306966173c2), OLFML3 (#50593011c1), TCTA (#148922970c1), TPP1 (#118582287c1), EMC1 (#22095330c3), IL7R (#28610150c2), EFHD1 (#237649043b1), PPIC (#45439319c2), NCEH1 (#226423949c2), CHST2 (#344925865c1), Actb (#6671509a1), and GAPDH (#378404907c1), ASCL1 (#190343011c1), BRN2 (#380254475c1), MAP2 (#87578393c1), MYT1L (#60498972c3), TUBB3 (#308235961c1), cMYC (#239582723c3), SOX2 (#325651854c3), KLF4 (#194248076c2), POU5F1(OCT4) (#4505967a1), NANOG (#153945815c1), DPPA2 (#239835766c1), DPPA4 (#144953902c1), REX1 (ZFP42) (#89179322c2).

### Retrovirus and lentivirus preparation

Large-scale retro-virus production was performed at the SBP Viral Vector Core Facility SBP as previously described 37. Briefly, for retrovirus preparation, pMX-MGT^76^, Retro-Gag-Pol, and pMD2.G plasmids were co-transfected into HEK-293T cells at a ratio of 3:2:1. For lentivirus preparation, lentivector DNA plasmids (Addgene plasmids # 27150, Tet-O-FUW-Ascl1; 27151, Tet-O-FUW-Brn2; 27152, Tet-O-FUW-Myt1l; 20342, 51543, FUW-tetO-hOKMS, FUW-M2rtTA, pLKO.1 shAtf7ip TRCN0000374251, pLKO.1 shJunb TRCN0000232241, pLKO.1 shSp7 TRCN000082147, pLKO.1 shZfp207 TRCN0000225905 were individually co-transected with pCMVDR8.74 and pMD2.G into HEK-293T cells using the calcium phosphate method. UltraCulture serum-free medium (Lonza) supplemented with 1 mM L-glutamine (Life Technologies) was used to re-feed transfected cells, and the supernatant was collected every 24 h from day 2 to day 4 after transfection. Viral supernatants were pooled, passed through a 0.22-μm-pore filter, concentrated, and purified by 20% sucrose gradient ultracentrifugation at 21,000 rpm for 2 h at 4°C. The pellet containing concentrated viral particles was resuspended in PBS, aliquoted, and kept at −80°C.

### Calcium handling assay

The calcium assay was performed on day 30 (HDFs) or day 20 (HAECs) after MGT overexpression. The assay was performed using the labeling protocol as previously described^77^. Briefly, 50% of the cell culture supernatant was replaced with a 2X calcium dye solution consisting of Fluo-4 NW dye (Invitrogen, F36206), 1.25 mM probenecid F-127 (Invitrogen), and 100 μg/mL Hoescht 33258 (Invitrogen, H3569, in water) diluted in warm Tyrode’s solution (Sigma), and the cells were incubated at 37°C for 45 min. The cells were then washed 4 times with fresh pre-warmed Tyrode’s solution and automatically imaged with an ImageXpress Micro XLS microscope (Molecular Devices) at an acquisition frequency of 100 Hz for a duration of 5 s with excitation 485/20 nm and emission 525/30 nm filters. A single image of Hoescht fluorescence was acquired before the time series. Fluorescence quantification over time and single-cell trace analysis were automatically performed using custom software packages developed by Molecular Devices and the Colas laboratory.

### RNA-seq and data analysis

HDFs were added to 384-well plates at 10^3^ cells/well and transfected with siRNAs (siATF7IP, siJUNB, siSP7, siZNF207 individually or in combination) in iCRM. The next day (day 0), the cells were transduced with 1 μL/well mouse MGT retrovirus diluted in iCRM. On day 2, cells were collected and RNA was extracted using TRIzol. Cells were pooled from 16 wells per sample, and two biological replicates per condition were analyzed. Library preparation was performed by Novogene using their in-house preparation protocol. Briefly, mRNA was enriched using oligo (dT) beads and fragmented randomly using a fragmentation buffer. cDNA was generated from mRNA template using a random hexamer primer followed by second-strand synthesis. Terminal repair, A ligation, and sequencing adaptor ligation were then performed. The final libraries were generated through size selection and PCR enrichment and sequenced as 2×150bp on a HiSeq2500 Sequencer (Illumina). Samples were sequenced to an approximate depth of 35–40 million reads per sample. Raw sequencing reads were trimmed using Trimmomatic (0.36) with a minimum quality threshold of 35 and minimum length of 36^78^. Processed reads were mapped to the hg38 reference genome using HISAT2 (2.0.4)^79^. Counts were then assembled using Subread featureCounts (1.5.2)^80^. Differential gene expression was analyzed using the DESeq2 package (1.20)^81^ in R. Genes were defined as differentially expressed if the adjusted p value was <0.05 after correction for multiple testing using the Benjamini–Hochberg method. Gene Ontology (GO) analysis was performed using PANTHER version 12.0 classification^82,83^.

### Single-cell ATAC-seq (scATAC-seq)

scATAC-seq experiments were performed with control (untreated) HDFs, MGT+siControl HDFs, and MGT+siAJSZ HDFs. HDFs were added to 384-well plates at 2.5×10^3^ cells/well in iCRM and transfected with siRNAs. The next day (day 0), cells were transduced mouse MGT retrovirus and collected 2 days later using trypsin-EDTA. Cells from 40 wells were pooled to obtain 2×10^5^ cells per sample, with two biological replicates per condition. Cells were washed with FCM, centrifuged in conical tubes, and the pellets were frozen in Freezing Medium (DMEM, 10% DMSO, 20% FBS), transferred to cryotubes, and placed in Mr. Frosty containers (Thermo Fisher) at −80°C.

Samples were processed for scATAC-seq at UCSD CMME. Samples were processed for scATAC-seq at UCSD CMME using 10x Genomics and sequenced on a NovaSeq 6000 at a depth of 20-25k read pairs per nucleus. Cell Ranger-ATAC (1.1.0) pipeline was used to filter and align reads, count barcodes, identify transposase cut sites, detect chromatin peaks, prepare t-SNE dimensionality reduction plots, and compare differential accessibility between clusters. The Cell Ranger-ATAC pipeline uses the following tools: cutadapt, BWA-MEM, SAMtools tabix, and bedtools. Further differential accessibility analysis was performed using DiffBind (2.12.1) and custom R scripts and visualized with ggplot2. Tracks were visualized using Integrative Genome Viewer 2.8.12. All scripts for this analysis are available on GitHub [https://github.com/smurph50].

### Chromatin immunoprecipitation-seq (ChIP-seq)

ChIP-seq experiments were performed with 10^8^ HDFs per replicate using a SimpleChip Plus sonication chromatin IP kit (Cell Signaling Technology, 56383) according to the manufacturer directions. In brief, HDFs were grown to 80% confluency (10^7^/sample) and then crosslinked with 1% formaldehyde (Sigma, F8775) in PBS at room temperature for 20 min with occasionally stirring. The crosslinking reaction was quenched by addition of 0.125 M glycine for 10 min, and chromatin was fragmented for 25 min using a Bioruptor Pico sonicator (Diagenode) to an average DNA fragment length of 200–500 bp. DNA was quantified with Qubit (Invitrogen, Q32854). Samples equivalent to 100 μg of DNA were incubated overnight at 4°C with 4 μg of rabbit polyclonal anti-ATF7IP (Invitrogen, PA5-54811), rabbit monoclonal anti-JUNB (C37F9) (Cell Signaling Technology, 3753S), rabbit anti-SP7/OSTERIX (Abcam, Ab22552), or rabbit polyclonal anti-ZNF207 (Bethyl laboratories, A305-814AM). Rabbit IgG (Cell Signaling Technology, 2729) was used as a negative control. Immunocomplexes were captured by rotation with protein G-coupled magnetic beads (Cell Signaling Technologies, 9006) for 2 h at 4°C, and immuno-precipitated genomic DNA was collected by incubation with 50 μL elution buffer. Library preparation and sequencing of immunoprecipitated and input DNA was performed by UCSD IGM core facility. Raw reads were mapped to GRCh38 using Bowtie2 (2.3.5). Since each sample was run across two lanes, SAM files were merged using Picard (2.20.5). MACS2 (2.1.1) was used to call narrow peaks relative to input with a q value cutoff of 0.01. Peaks were annotated with Homer (4.10.4) and motifs were analyzed using MEME-ChIP (5.1.1). BigWig files were generated using Deeptools (2.2) bamCoverage. Tracks were visualized with Fluff (biofluff 3.0.3). Gene ontology biological process terms were found with PANTHER GO and overlap analyses were performed using custom R scripts with venneuler and ggplot2 packages. Bedtools (v2.29.2) was used for genomic comparisons and combining ChIP-seq and scATAC-seq data. HOMER was used to find motifs with a scrambled background.

### Mouse MI model

Experiments were performed in 10-to 12-week-old randomly allocated male and female mice (Jackson lab, strain 000664). To generate the mouse MI model, mice were intubated and anesthetized with isoflurane gas. The chest cavity was exposed by cutting the intercostal muscle and then the left coronary artery was ligated with a 5-0 silk suture.

### Echo-guided retro/lenti-viruses injection

3 days after MI, 6 μL of virus-containing solution (PBS) was injected into the boundary between the infarct and border zone at one site with a 32-gauge needle using echo-guided visualization as described in 62. The mouse surgeon was blinded to the study and mortality after MI was < 10%.

### Echocardiography

Cardiac function was analyzed with transthoracic echocardiography (Visual Sonics, Vevo 2100) at 2 weeks, and 4 weeks after MI (n = 7 per group). Mice were anesthetized with low dose isoflurane for echocardiographic examination. Two-dimensional targeted M-mode traces were obtained at the papillary muscle level. Left ventricular internal diameter during diastole (LVDd) and left ventricular internal diameter during systole (LVDs) were measured in at least three consecutive cardiac cycles. EF and FS were calculated with the Teichholtz formula.

### Statistical analysis

All statistical analyses were performed using Prism version 8.0 (GraphPad Software, San Diego CA, USA). Data are presented as the mean ± SEM unless noted. Statistical significance was analyzed by unpaired Student’s t-test or one-way ANOVA. P values of <0.05 were considered significant.

## Acknowledgements

We thank Kirsten Jepsen (UCSD IGM Genomic Core Facility) for assistance with the ChIP-seq experiments; Allen Wang and Sebastian Preissl (UCSD CMME) for assistance with the scATAC-seq experiments; Luca Caputo, Haley Vaseghi, and Li Wang for kindly sharing reagents and instruments; and Sean Spiering, Eleanor Kim, and Josiah Punay for excellent technical support.

---

**Supplemental Figure 1.**
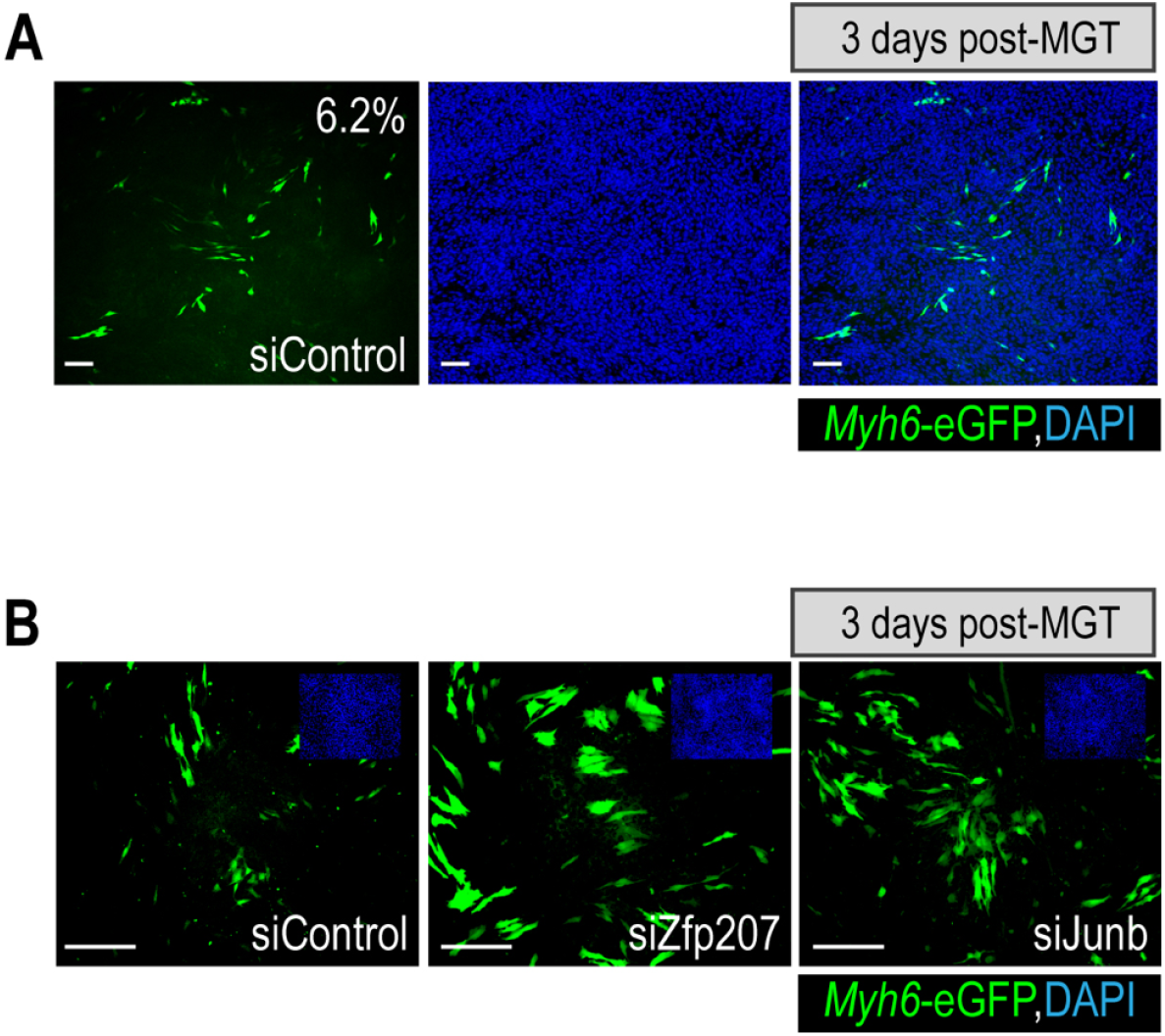
**(A and B)** Representative images of siControl-transfected with average reprogramming efficiency quantification (6.2% of Myh6-eGFP+ cells) (A) and siControl-, siZfp207-, or siJunb-transfected (B) iMGT-MEFs 3 days after MGT overexpression. The cardiac marker Myh6 is shown in green, and cell nuclei are stained blue (DAPI). Scale bars: 50 μm.

**Supplemental Figure 2.**
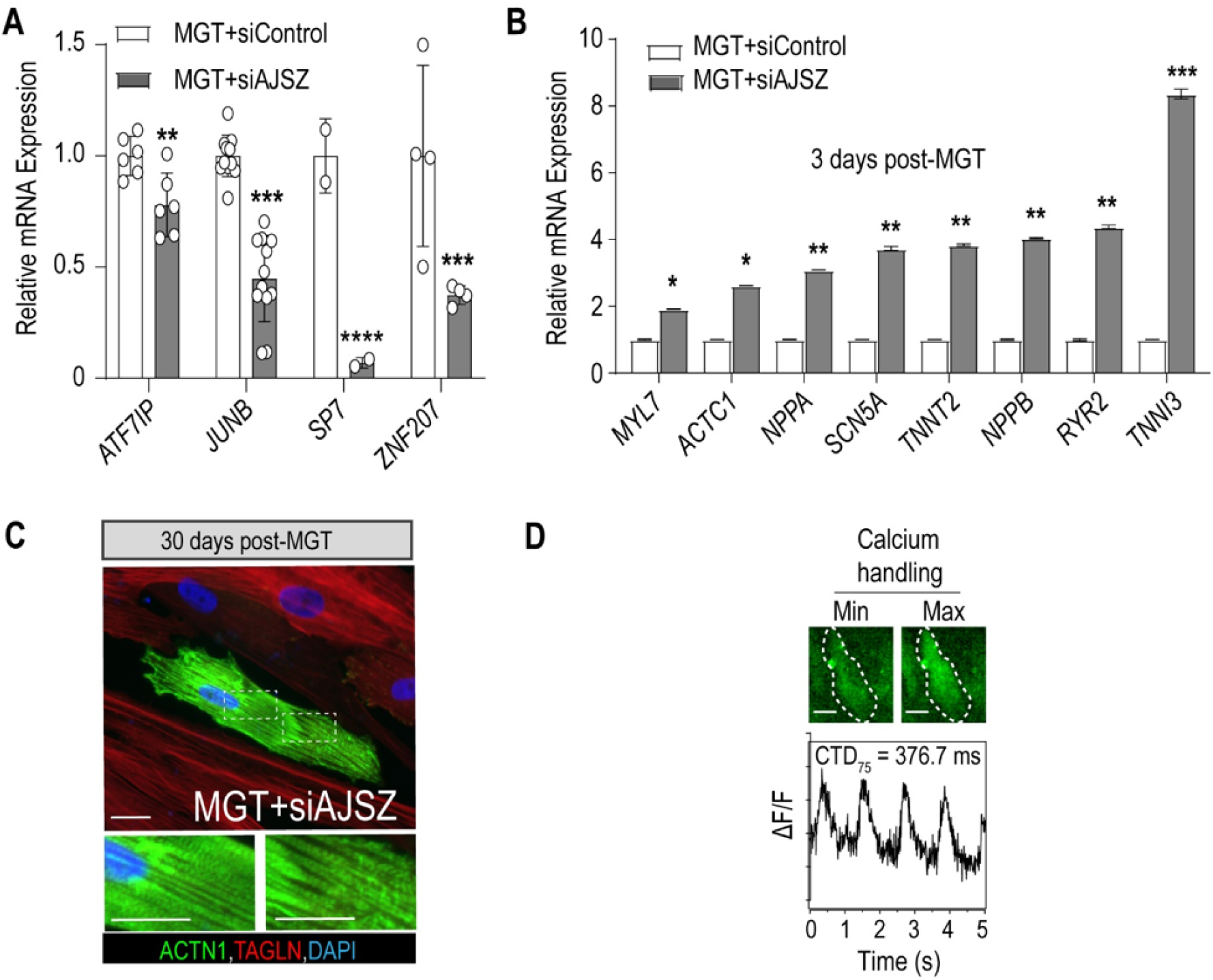
**(A)** qRT-PCR analysis of AJSZ expression in siCTR- or siAJSZ-transfected HDFs 3 days after mMGT overexpression. Data were normalized to the MGT+siCTR cells. **(B)** qRT-PCR analysis of the indicated cardiac gene expression in siCTR- or siAJSZ-transfected HDFs 3 days after mMGT overexpression. **(C)** Immunostaining of ACTN1 and TAGLN in siAJSZ-transfected HDFs analyzed 30 days after MGT overexpression. Lower panels show that some ACTN1+ cells have lost TAGLN staining and show striations. **(D**) Fluorescence-based (Fluo-4) quantification of calcium handling in siAJSZ-transfected iMGT-MEF cells 30 days after mMGT overexpression. Scale bars: 50 μm. Student’s t-test. *p<0.05, **p<0.01, p***<0.001, ****p<0.0001.

**Supplemental Figure 3.**
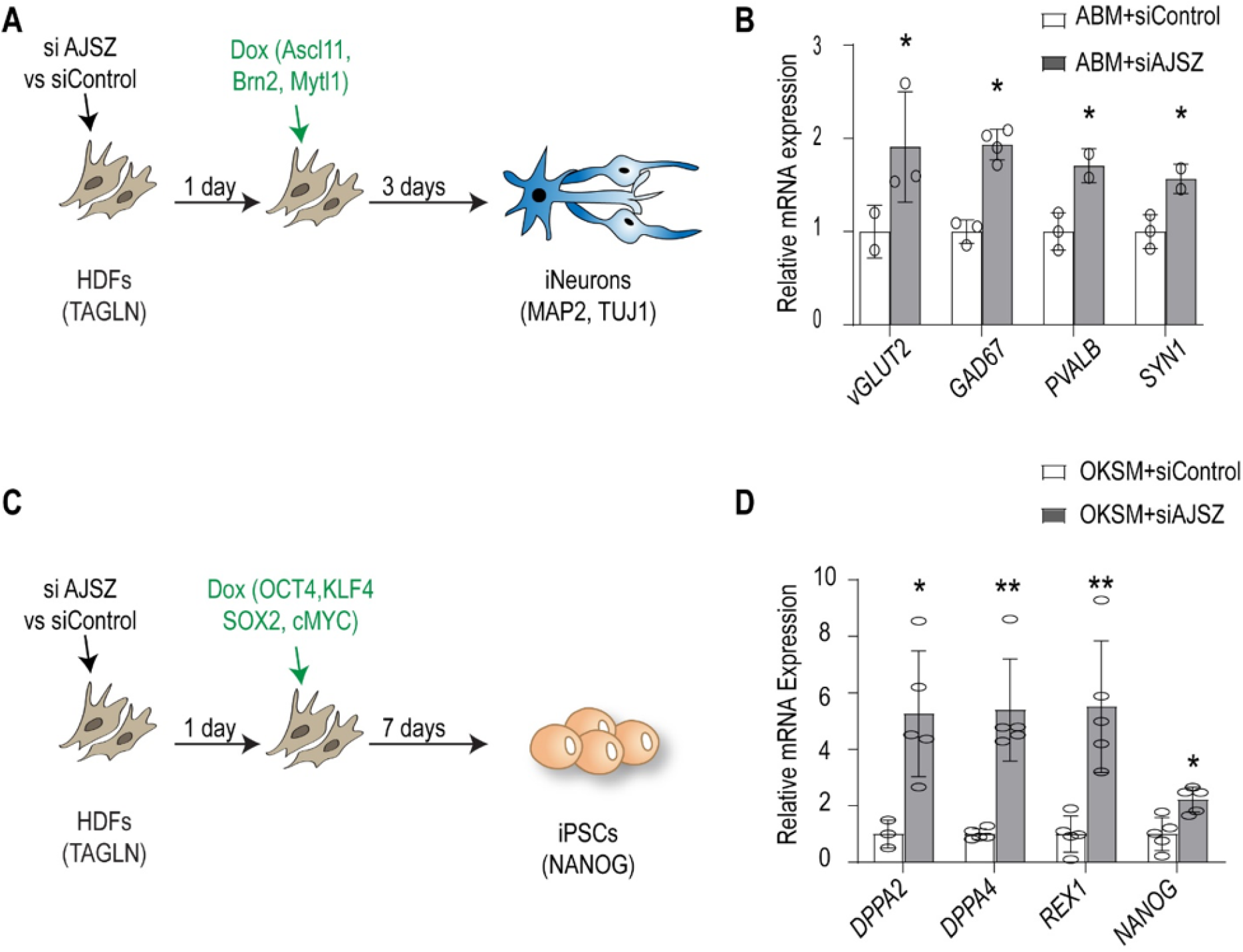
**(A)** Schematic showing the experimental set-up for direct neuronal reprogramming of HDFs with ABM. **(B)** qRT-PCR analysis of neuronal markers in siRNA-transfected HDFs, 3 days after induction of neuronal reprogramming with ABM. **(C)** Schematic showing the experimental set-up for reprogramming of HDFs into iPSCs with OKSM. **(D)** qRT-PCR analysis of pluripotent markers in siControl- and siAJSZ-transfected HDFs on day 7 after OKSM overexpression. Student’s t-test. *p<0.05, **p<0.01.

**Supplemental Figure 4.**
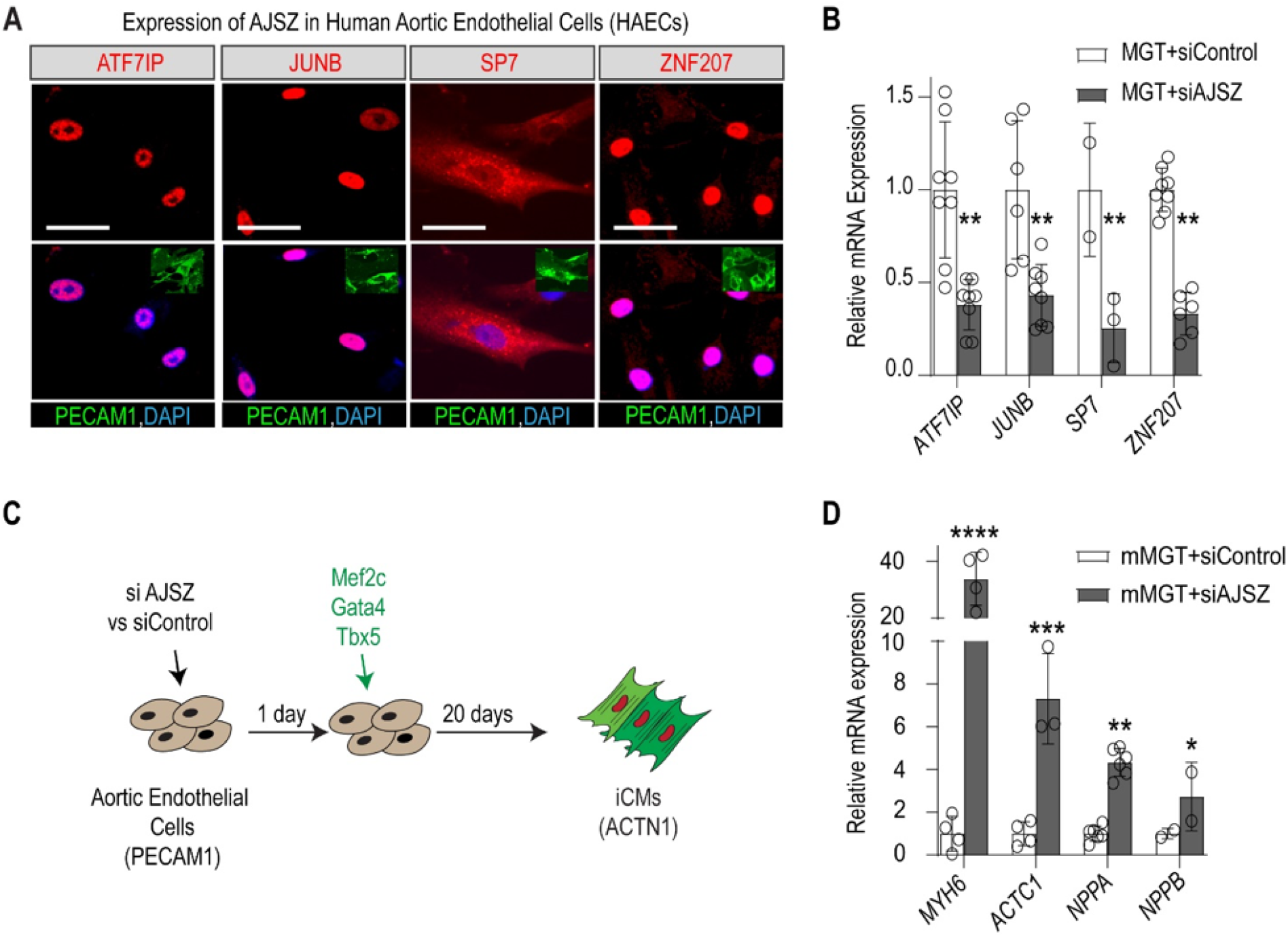
**(A)** Immunostaining of AJSZ (red) and endothelial marker PECAM1 (green) in HAECs. Nuclei are stained with DAPI (blue, top left insets). Scale bars: 50 μm. **(B)** qRT-PCR analysis of AJSZ expression in siControl- and siAJSZ-transfected HAECs on day 2 after mMGT overexpression. **(C)** Schematic depicting direct cardiac reprogramming assay in HAECs. **(D)** qRT-PCR of cardiac genes in siControl- and siAJSZ-transfected HAECs on day 3 after mMGT overexpression. Student’s t-test. *p<0.05, **p<0.01, ***p<0.001, ****p<0.0001. Scale bars: 50 μm.

**Supplemental Figure 5.**
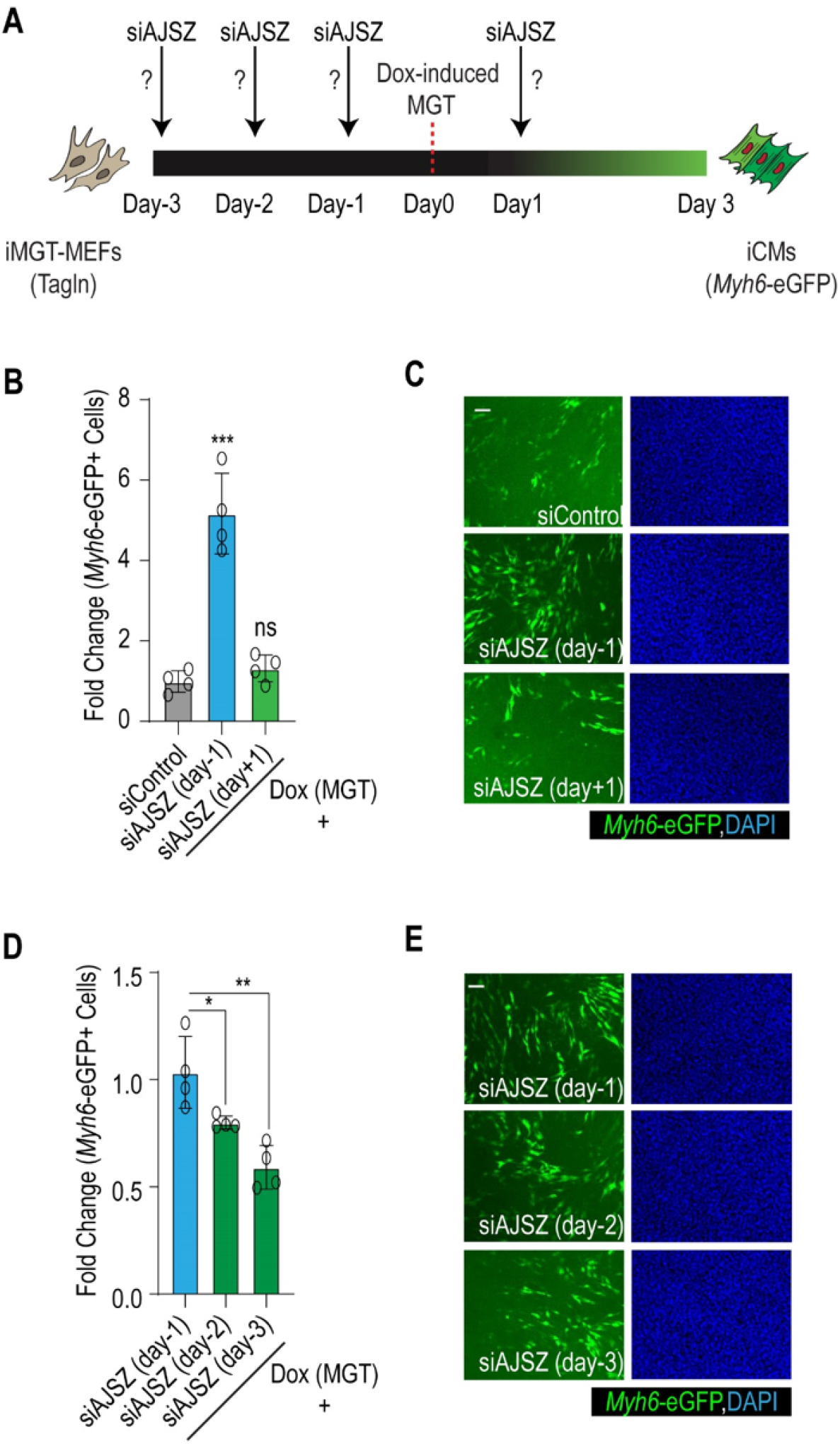
**(A)** Experimental strategy to determine optimal AJSZ KD timing, to elicit maximal CR efficiency in iMGT-MEFs assay. **(B, C)** Quantification (B) and representative images (C) of reprogramming efficiency in response to AJSZ KD one day prior or one day after MGT overexpression. **(D, E)** Quantification (D) and representative images (E) of reprogramming efficiency in response to AJSZ KD 1, 2, or 3 days before induction of MGT. two-way ANOVA, *p<0.05, **p<0.01, ***p<0.001. Scale bars: 50 μm.

**Supplemental Figure 6.**
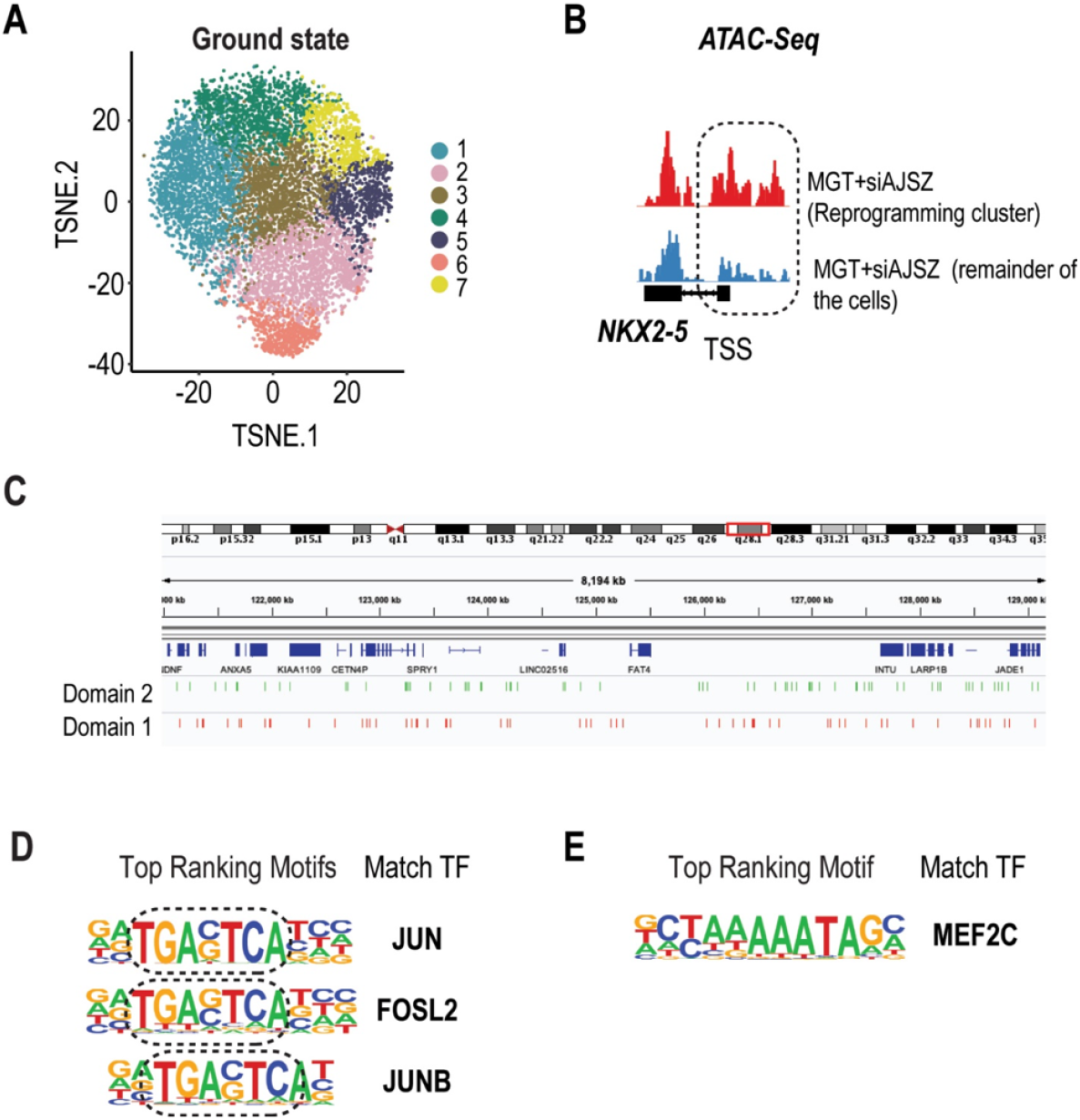
**(A)** t-SNE visualization of cell clusters in HDFs at ground state using scATAC-seq. **(B**) ATAC-seq track for NKX2.5 at the TSS regions in HDFs in reprogramming cluster as compared to the remainder of the cells. **(C)** Example of *domain 1* and *domain 2* distribution across ~8MB at region 4 q28.1. *domain 1* and 2 consist in short stretches of DA chromatin evenly distributed across this region. (**D)** Top 3 motifs enriched in *domain 1* are AP-1 TF motifs. **(E)** Second most motif enriched in *domain 2* corresponds to reprogramming TF MEF2C putative DNA binding site.

**Supplemental Figure 7.**
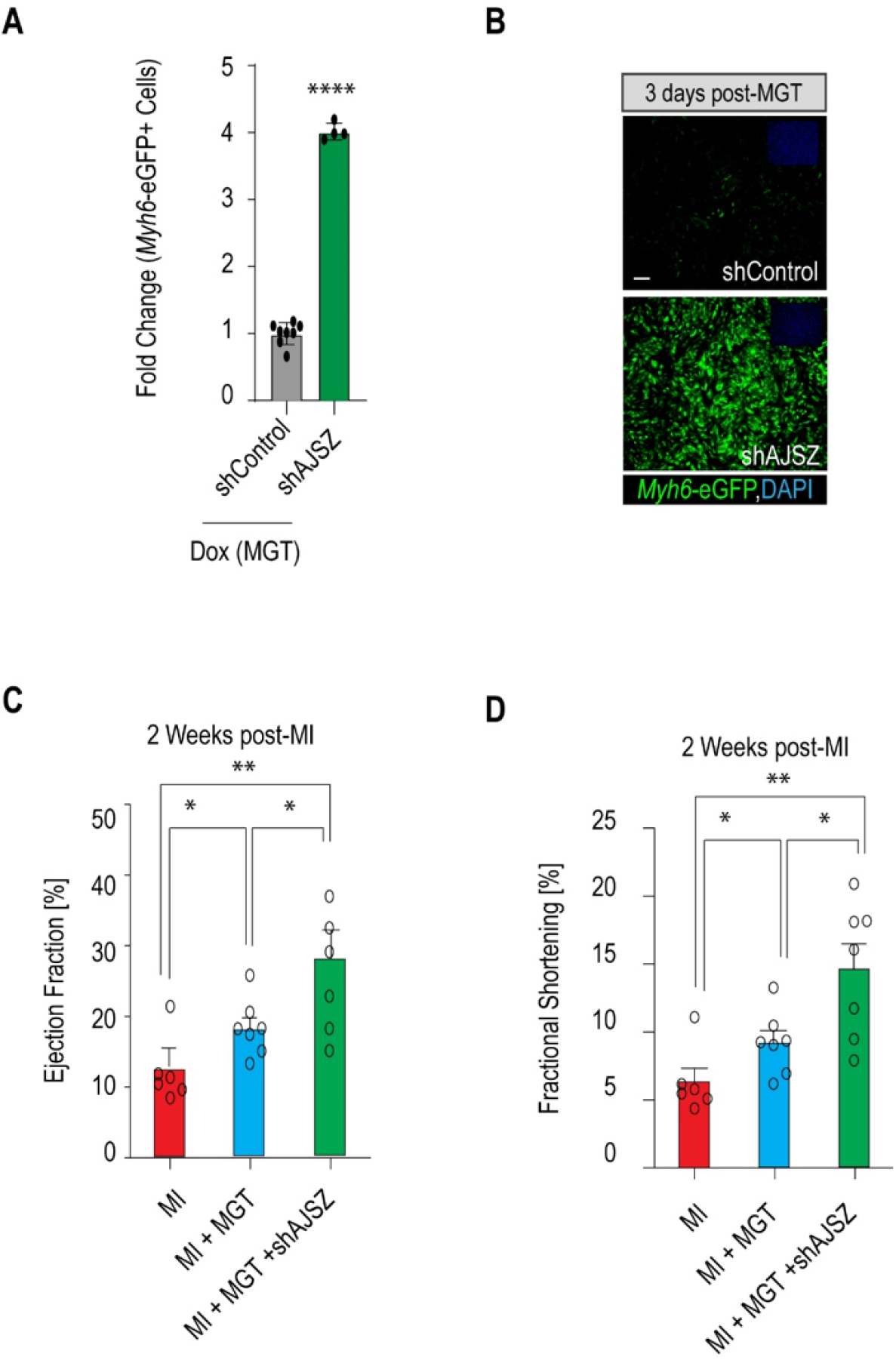
**(A)** shRNA-mediated KD of AJSZ enhances CR efficiency in iMGT-MEFs as compared to control shRNA. **(B)** Representative images of shAJSZ- and shControl-infected iMGT-MEFs on day 3 after MGT induction. **(C)** Quantification of the percentage of Zfp207+ cells in heart section in MGT and MGT+shAJSZ. **(D)** Immunostaining of Zfp207 (red) and DAPI (blue) in heart sections from MGT or MGT+shAJSZ injected mice, showing that injection that shAJSZ induces a significant decrease of the percentage of Zfp207 expressing cells as compared to MGT. *p<0.05, **p<0.01, ****p<0.0001. Scale bars: 50 μm.

**Supplemental Figure 8.**
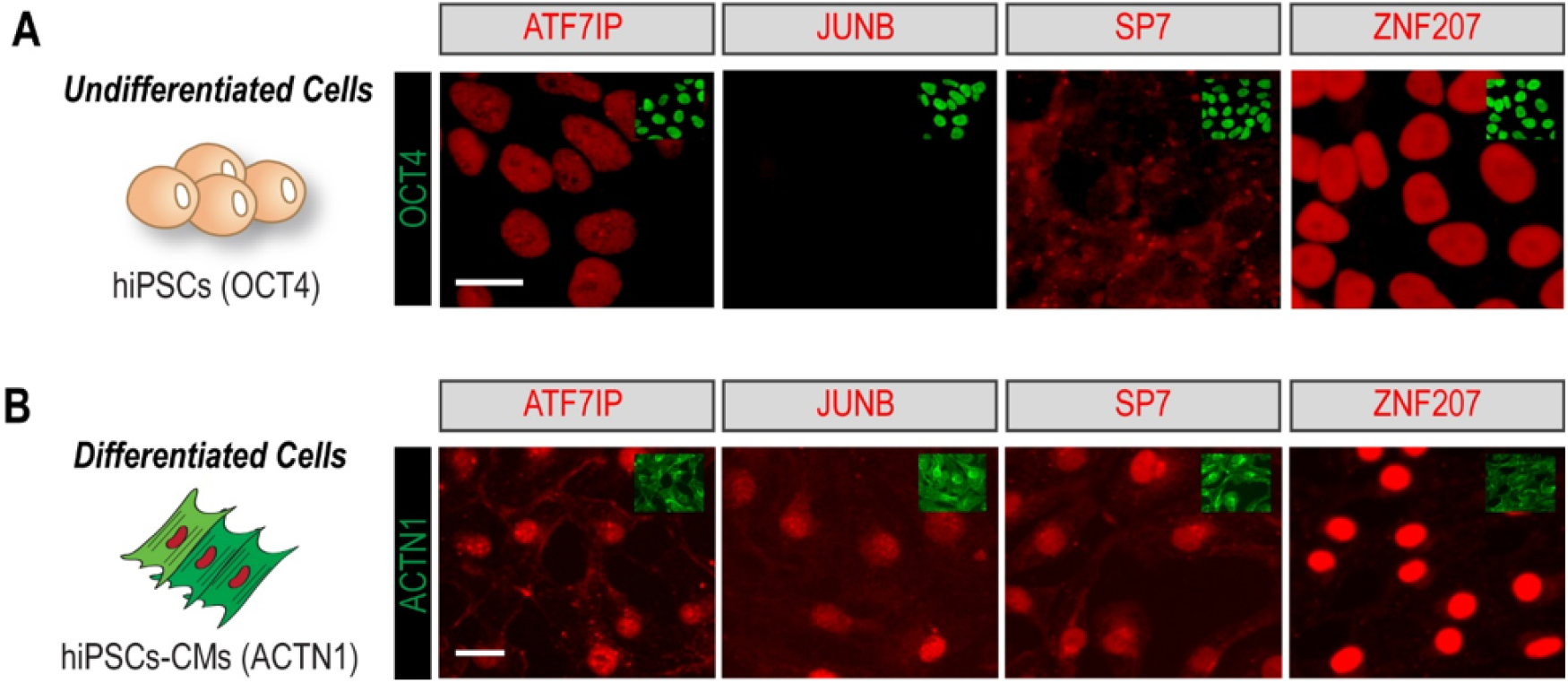
**(A)** Immunostaining of AJSZ (red) and pluripotency marker OCT4 (green) in hiPSCs. **(B)** Immunostaining of AJSZ (red) and marker of cardiac differentiation ACTN1 (green). Nuclei are stained with DAPI (blue, top left insets). Scale bars: 30 μm.

